# Air Pollution, Early Adversity, and Amygdala: Environmental Correlates of Psychopathology in Preadolescents

**DOI:** 10.64898/2026.08.31.748438

**Authors:** Amanda C. Del Giacco, Michael A. Rosario, Carlos Cardenas-Iniguez, Nitya Chawla, Alan Wen, Kirthana Sukumaran, L. Nate Overholtzer, Jiu-Chiuan Chen, Benjamin B. Lahey, Tyler M. Moore, Megan M. Herting

## Abstract

Early life adversity (ELA) and ambient fine particulate matter (PM_2.5_) are hypothesized to be environmental risk factors for altered brain structure and psychopathology, but their unique and interactive effects in childhood remain unclear. This study examines the interactive associations of ELA and annual average residential PM_2.5_ exposure (total mass and 15 components) on total amygdala and basolateral amygdala subregion volumes and psychopathology symptoms in a subset of children (N=3,601, 45% assigned female at birth, 9-10 years) from the Adolescent Brain Cognitive Development Study. Linear mixed-effects models, adjusting for sociodemographic factors, co-pollutants, and neuroimaging covariates, showed that ELA was associated with greater bifactor model-defined general, specific internalizing, and specific externalizing symptoms of psychopathology. PM_2.5_ moderated ELA associations with specific externalizing symptoms, with greater symptoms in those exposed to higher ELA and higher PM_2.5_ exposures. Among youth with higher ELA, smaller basolateral paralaminar volumes were linked to greater general and specific externalizing symptoms. These findings underscore the importance of considering psychosocial and physical environmental co-exposures when identifying children at risk for psychopathology.

## 1. Introduction

Childhood and the transition to adolescence represent sensitive periods for the development of psychiatric disorders (McGrath et al. 2023; Paus, Keshavan, and Giedd 2008). Early life adversity (ELA) and ambient air pollution are increasingly recognized as risk factors for altered brain development and mental health disorders (Hobbs et al. 2025; Morrel, Dong, et al. 2025; Nelson, Sullivan, and Valdes 2025; Wade, Wright, and Finegold 2022). Studies investigating their interactive effects suggest that experiencing both a psychosocial and physical environment stressor together may magnify or alter the combined impact on brain development (de Jesus et al. 2025; Lubczyńska et al. 2021; Rakesh, Zalesky, and Whittle 2022) and mental health (Pagliaccio et al. 2020; Vargas, McLaughlin, and Rakesh 2025; Zundel et al. 2022). However, others showed minimal additional associations of ambient air pollution for children experiencing high-stress environments (Miller et al. 2022). These divergent findings map onto two competing frameworks. Under a ‘double hit’ framework, one toxic exposure may sensitize the developing stress system, such that a subsequent environmental exposure produces disproportionately greater susceptibility for psychopathology than a single exposure alone (Clougherty and Kubzansky 2009). In contrast, under a ‘saturation’ model, a toxic exposure has engaged much of the system’s adaptive capacity, leaving less room for an additional exposure to further impact brain structure and behavior (Clougherty et al. 2006). To better understand mental health pathogenesis and hypotheses regarding how these environmental risk factors might become biologically embedded, it is crucial to examine how these contextual drivers interact in their associations with individual differences in mental health disorders and their underlying neural substrates.

ELA is characterized by disadvantageous experiences during early life, including unstable households, physical and emotional abuse and neglect, witnessing violence, and parental substance abuse or mental illness (Breslin et al. 2025; Duffy, McLaughlin, and Green 2018). These experiences occur frequently, with approximately 80% of adolescents reporting at least one ELA (McLaughlin et al. 2012; Swedo et al. 2024). Retrospective studies in adults have consistently linked ELA to greater psychopathology symptoms, as well as elevated rates of depression, anxiety, and substance misuse (Dosanjh et al. 2025). Similarly, in children, greater ELA has been associated with increased internalizing and externalizing symptoms (Kim and Cicchetti 2010; Vachon et al. 2015; Wade et al. 2022) and behavioral problems (Choi, Wang, and Jackson 2019).

Exposure to fine particulate matter (PM_2.5_, with an aerodynamic diameter ≤2.5 μm) has similarly been linked to poor mental health outcomes (Mazahir, Shukla, and Albastaki 2025; Zundel et al. 2022). Children exposed to higher levels of PM_2.5_ had greater emotional and behavioral difficulties (Ahmed et al. 2022), and adolescents in the highest PM_2.5_ exposure range show elevated psychopathology symptoms (Reuben et al. 2021). However, findings are not consistent. One large cohort study found no association between PM_2.5_ exposure and emotional and conduct problems in children aged 11 to 12 years (Bradley et al. 2024), and another study did not demonstrate a relationship between PM_2.5_ exposure and emotional and behavioral problems in children over time (Campbell et al. 2024). A missing component of these studies is the lack of research investigating how ELA and air pollution exposure may together relate to mental health outcomes. Notably, children living in disadvantaged neighborhoods are disproportionately exposed to both higher rates of ELA and higher concentrations of harmful air pollutants (Merrick et al. 2018; Wodtke et al. 2022). This raises the possibility that these exposures frequently co-occur and may interact, consistent with either the ‘double hit’ or ‘saturation’ frameworks, to shape mental health risk. Additionally, not all children with elevated exposures to risk factors develop mental health disorders, underscoring the need to better understand the individual and combined pathways through which ELA and PM_2.5_ might increase psychopathology risk. Given that ELA and air pollution have each been linked to a broad, overlapping range of psychiatric symptoms and diagnoses rather than any single disorder (Mazahir et al. 2025; McLaughlin, Weissman, and Bitrán 2019; Wade et al. 2022; Waters and Gould 2022), capturing their interactive effects may require moving beyond category-specific outcomes. That is, researchers may need to turn toward transdiagnostic behavioral and brain biomarkers (i.e., looking at mental health factors that cut across different traditional diagnostic categories) capable of indexing shared liability across disorders.

The amygdala represents a focal brain region biologically implicated in stress and emotion processing, given its central anatomical location and functional integration with many cortical and subcortical regions (Joëls and Baram 2009; Phelps and LeDoux 2005). The amygdala also undergoes significant developmental changes from childhood through adolescence (Herting et al. 2018), making late childhood an optimal time to understand how psychosocial and physical environmental exposures might impact its development. Furthermore, the amygdala has been implicated in multiple mental health disorders (e.g., anxiety, depression, alcohol misuse, posttraumatic stress disorder) (Cao et al. 2026; Gilpin, Herman, and Roberto 2015; Oshri et al. 2024; Sharp 2017; Shin, Rauch, and Pitman 2006), positioning it as a candidate transdiagnostic neural substrate for shared risk of multiple disorders (Cao et al. 2026; Sambuco, Bradley, and Lang 2023). Structural differences in the amygdala have also been linked to both ELA (Hanson and Nacewicz 2021) and specific components within PM_2.5_ exposure (Guxens et al. 2018; Morrel, Overholtzer, et al. 2025). However, these literatures have developed largely in parallel, and minimal work has examined the interactive effects of ELA and air pollution on the emotion-processing hub of the amygdala. This gap is compounded by the fact that the amygdala is not a unitary structure, as its subregions demonstrate distinct functional profiles (Bzdok et al. 2013; deCampo and Fudge 2012), and volumetric developmental variations (Fish et al. 2020) that may be obscured when looking at the entire structure as a whole. The basolateral complex of the amygdala (BLA) in particular contributes to conditioned fear and stress responses (Keding and Herringa 2016; Sepahvand et al. 2023) and an individual’s susceptibility to psychopathology (Prager et al. 2016; Sharp 2017). A small number of studies have examined amygdala subregions in relation to a single environmental exposure. For example, a recent paper from our group linked amygdala subregions to specific components of PM_2.5_ total mass (Morrel, Overholtzer, et al. 2025), but this study did not examine behavioral or psychopathology outcomes. Thus, a critical gap in knowledge is examining whether ELA independently and/or jointly with PM_2.5_ exposure relates to BLA subregion volumes and transdiagnostic mental health symptoms. Accordingly, the BLA represents a biologically plausible neural substrate through which ELA and PM_2.5_ exposure may exert combined effects on mental health and behavioral outcomes.

The present study investigates the unique and interactive associations of ELA and annual average PM_2.5_ exposure with amygdala structure and psychopathology in preadolescents (**Figure 1**). Using cross-sectional data from the Adolescent Brain Cognitive Development^SM^ (ABCD) Study baseline assessment (Garavan et al. 2018; Karcher and Barch 2021; Volkow et al. 2018), we tested three independent moderation models. First, we examined the independent and interactive associations of ELA and PM_2.5_ exposure with (1) total amygdala and BLA subregion volumes and (2) psychopathology symptoms, indexed by a bifactor model capturing general (p), specific internalizing, and specific externalizing dimensions (Moore et al. 2020). Then, we integrated these components into a single model to determine how ELA, PM_2.5_ exposure, and BLA subregion volumes all relate to psychopathology symptoms, testing if ELA and PM_2.5_ moderated associations between BLA subregion volumes and mental health symptoms. Given prior evidence from our group that specific PM_2.5_ components (e.g., elemental carbon, organic carbon, nitrates) can show associations with brain and behavioral outcomes that diverge from those observed for PM_2.5_ total mass (Morrel, Overholtzer, et al. 2025; Rosario et al. 2026; Sukumaran et al. 2024, 2025), we also conducted exploratory analyses of a potential mixture effect of individual PM_2.5_ components, which may otherwise be obscured when only total mass is considered. Based on previous findings, we hypothesized that ELA would be associated with smaller BLA volumes (Hanson and Nacewicz 2021; Herzog et al. 2020; VanTieghem et al. 2021), and that these relationships would be stronger for youth exposed to higher levels of ambient air pollution (Margolis et al. 2022; Pagliaccio et al. 2020). Additionally, we hypothesized that ELA would be positively related to psychopathology (Callaghan and Tottenham 2016; Kessler et al. 2010; Nelson et al. 2025) and PM_2.5_ exposure would moderate this association (Cattarinussi et al. 2026; de la Rosa et al. 2024), in line with the ‘double hit’ hypothesis. Finally, we hypothesized that BLA volumes would be negatively related to psychopathology symptoms (Oshri et al. 2019; Ousdal et al. 2020), and that ELA and air pollution would moderate these associations, such that smaller BLA subregion volumes would be associated with higher psychopathology symptoms among youth with higher ELA and greater PM_2.5_ exposure.

**Fig. 1.**
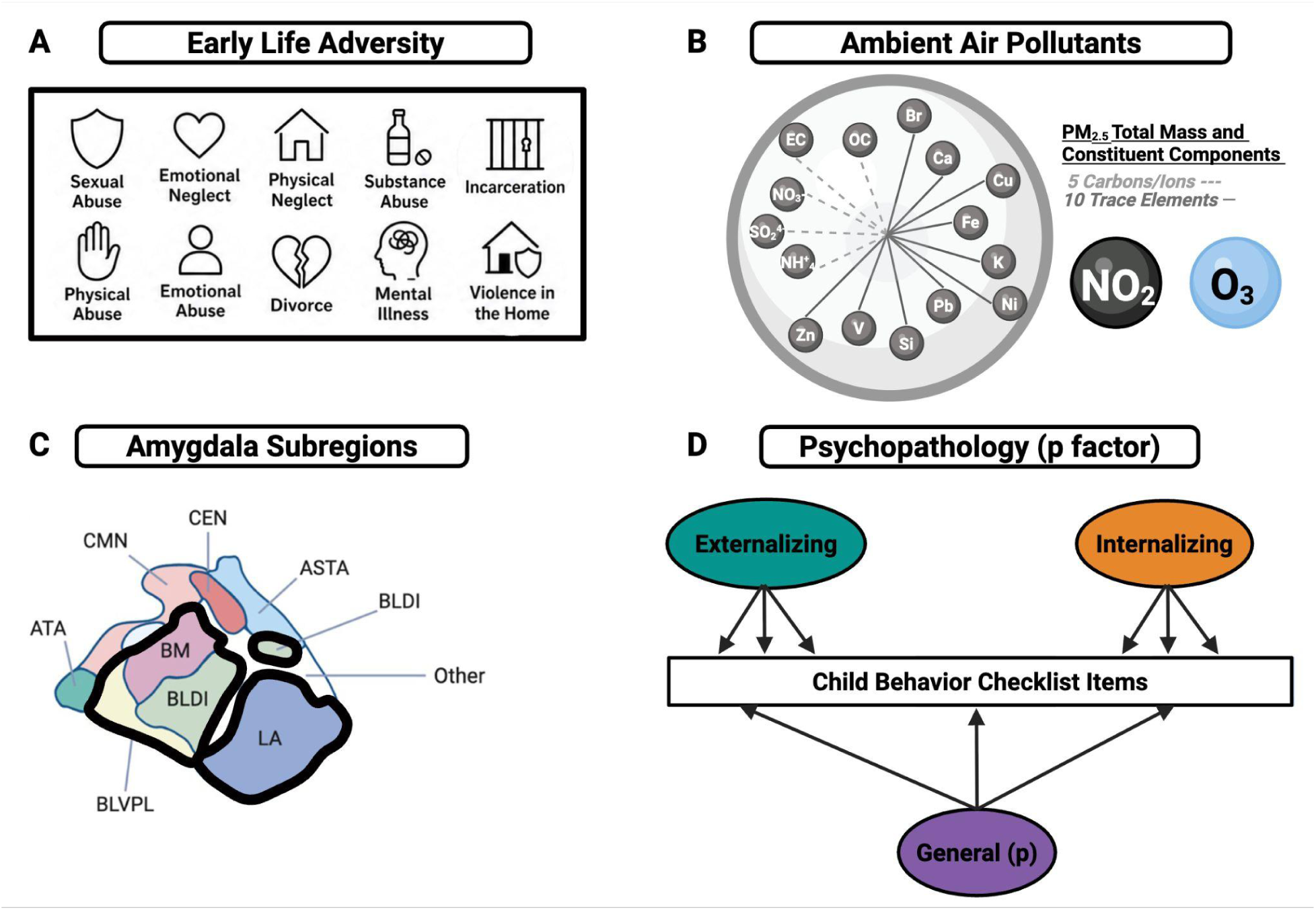
Study Variables and Measurement Framework. **A.** Early Life Adversity (ELA) was calculated as the total sum of 10 adverse experiences as determined in Breslin et al., 2025. **B.** Ambient air pollution included in our analytical models included annual average PM_2.5_ total mass and its constituent components, while adjusting for co-exposure for NO_2_, and O_3_. **C.** Total amygdala volume and amygdala subregions were defined using the CIT168 atlas, with *a priori* selection of four basolateral complexes of the amygdala (BLA) subregions for the current study (outlined in black), including: lateral nucleus (LA), dorsal and intermediate divisions of the basolateral nucleus (BLDI), ventral division of the basolateral nucleus and paralaminar nucleus (BLVPL), and basomedial nucleus (BM). **D.** Bifactor models of psychopathology were based on Moore et al., 2020 and included general (p), and specific internalizing, and specific externalizing symptom factors.

## 2. Methods

### 2.1 Study design and dataset

The current study used a subsample from the baseline visits from the larger ABCD^®^ Study (Garavan et al. 2018; Volkow et al. 2018). The ABCD Study is the largest 10-year study of childhood neurodevelopment, which enrolled over 11,800 children (ages 9 to 10 years at baseline) between 2016 and 2018 across 21 sites in the United States. Child exclusion criteria for the ABCD Study included: limited child English proficiency, premature birth, intellectual disability, major medical, sensory, or intellectual concerns, and MRI incompatibility (Palmer et al. 2022). Parents and/or guardians provided written informed consent, and the youth provided written assent. All study procedures were approved by the central Institutional Review Board (IRB) at the University of California, San Diego, and each study site obtained local IRB approval. Further details about the conception and recruitment for the ABCD Study have been reported previously (Garavan et al. 2018; Volkow et al. 2018).

The data used for this study included raw T1- and T2-weighted MRI scans (ABCD Release 3.0; https://nda.nih.gov/study.html?id=1042), the ELA construct (ABCD Release 5.1; https://nda.nih.gov/study.html?id=2313), and variables including air pollution, p-factor, and demographic measures (ABCD Release 6.1; https://www.nbdc-datahub.org/abcd-release-6-1). From these data, participants were selected for the current study if they were enrolled at an ABCD study site with a Siemens scanner (excluding GE and Philips scanners; see section 2.4 *Total Amygdala and BLA Volumes*), passed ABCD MRI imaging quality-control procedures, had no incidental radiological findings on MRI, and had complete data for the primary exposure variables. To meet statistical modeling assumptions and reduce nonindependence, we randomly selected one child from each family. The final analytical sample was N = 3,601, with additional selection criteria details in **Supplemental Figure 1**.

### 2.2 Early Life Adversity (ELA)

As previously published (Breslin et al. 2025; Brieant et al. 2023), the ELA construct was generated from a systematic review of ABCD consortium projects and aimed to align ABCD Study ELA domains with the well-established 10-item adverse childhood experiences (ACEs) questionnaire (Felitti et al. 1998). These ABCD-assessed domains included: experiences of physical abuse, physical neglect, emotional abuse, emotional neglect, sexual abuse, parental mental illness, parental substance use disorders, witnessing violence in the home, parental divorce, and caregiver incarceration. The endorsement of each of these ELAs was measured across nine different parent- and youth-reported assessments within the ABCD battery, including the computerized Kiddie Schedule of Affective Disorders and Schizophrenia (KSADS-COMP), Children’s Report of Parent Behavior Inventory, Life Events Scale, Family Environment Scale, Adult Self-Report of the Achenbach System of Empirically Based Assessment, PhenX Neighborhood Crime and Safety Scale, and family history and demographics questionnaires, and were all measured at baseline (Karcher and Barch 2021). Each ELA domain was scored (0 = “absent” or 1 = “present”), and an assignment of ‘1’ was given if either child or caregiver endorsed any of the assessment items for that domain. The ELA measure used in the current study examined the total ELA summary score, generated by summing across ELA domains (0-10 counts) at baseline, when children were ages 9-10 years old. Higher summary scores represented more adverse life events experienced by youth.

### 2.3 Ambient Air Pollution Exposure

Annual average ambient air pollution concentrations, including daily measures of PM_2.5_ (µg/m^3^), nitrogen dioxide (NO_2_; ppb), and daily 8-hour (maximum) ground-level ozone (O_3_; ppb), were estimated for the primary address of each child, as previously described (Cardenas-Iniguez et al. 2024; Fan et al. 2021). Specifically, hybrid spatiotemporal models combining land-use regression, satellite-based aerosol optical depth, weather data, and chemical transport were used to derive daily estimates at a 1-km^2^ resolution across the continental United States (Di et al. 2019; Requia et al. 2020). Similar machine learning models were used to estimate monthly estimates of 15 PM_2.5_ components at a 50-meter resolution, including 5 carbons and ions (ammonium (NH₄⁺), elemental carbon (EC), nitrate (NO_3_⁻), organic carbon (OC), and sulfate (SO₄^2^⁻)) and 10 trace elements (bromine (Br), calcium (Ca), copper (Cu), iron (Fe), lead (Pb), nickel (Ni), potassium (K), silicon (Si), vanadium (V), and zinc (Zn)) (Jin et al. 2022). Model performance for these exposures across the US is reported in **Supplemental Table 1** (Sukumaran et al. 2025). These exposure estimates for PM_2.5_, NO_2_, O_3_, and PM_2.5_ components were averaged for the 2016 calendar year to correspond to an annual average exposure during the baseline enrollment period of the ABCD Study when the children were 9-10 years old. These estimates were then assigned to the primary residential address of each child provided by the caregiver.

### 2.4 Total Amygdala and BLA Volumes

As previously published, a harmonized 3T MRI data protocol was utilized to collect T1-weighted (T1w) and T2-weighted images across all ABCD Study sites (Casey et al. 2018). To segment the amygdala into subregions, we used the CIT168 atlas, a high-resolution *in vivo* probabilistic atlas of human amygdala subregions (Tyszka and Pauli 2016). The CIT168 atlas construction, validation, comparison with other atlases, and individual difference estimates have been reported previously, as well as specifics regarding each segmented subregion and its success in applying the subregion estimates in pediatric samples (Azad et al. 2021; Campbell et al. 2021, 2022; Kim et al. 2020). Because the CIT168 atlas was developed and validated using Siemens MRI data (Pauli, Nili, and Tyszka 2018; Tyszka and Pauli 2016), and the ABCD Study has reported large differences between scanner manufacturers, we included only participants at the 13 ABCD sites with Siemens scanners. For each participant, *in vivo* probabilistic volumes for nine amygdala subregions per hemisphere were quantified as previously described (Morrel, Overholtzer, et al. 2025; Overholtzer et al. 2025). This included application of a B-spline bivariate symmetric normalization diffeomorphic registration algorithm registered to the CIT168 atlas, after which the inverse diffeomorphism was applied to map CIT168 probabilistic atlas labels into individual space, and probabilistic label volumes were calculated via weighted summation of label probability over total volume. Following prior work (Campbell et al. 2021, 2022; Overholtzer et al. 2025), we included only participants whose amygdala segmentations had a contrast-to-noise ratio (CNR) ≥ 1.0 for all four measurements (left and right segmentations on both T1- and T2-weighted images).

Given the importance of the BLA in emotional processing and risk for psychopathology (Becker et al. 2023; Roozendaal, McEwen, and Chattarji 2009; Vyas et al. 2020), we chose *a priori* to examine the four BLA CIT168 nuclei, including lateral amygdala subregions (lateral nucleus (LA), dorsal and intermediate divisions of the basolateral nucleus (BLDI), ventral division of the basolateral nucleus and paralaminar region (BLVPL), and basomedial region (BM)). Each region of interest (ROI) was collapsed and summed across left and right hemispheres and then z-scored for standard comparisons. To account for head size differences and handedness-related differences, we adjusted brain volumes for total intracranial volume (ICV) and handedness, respectively.

### 2.5 Psychopathology Symptoms

As previously published, we implemented the bifactor model using the caregiver report of the Child Behavior Checklist (CBCL) (Achenbach and Rescorla 2001) within the ABCD Study sample (Moore et al. 2020). In this bifactor model, each CBCL mental health item loads simultaneously on a general factor (p-factor) and one orthogonal specific factor. This technique captures shared variance across all symptoms (general psychopathology) as well as unique variance to each specific symptom domain (e.g., internalizing, externalizing) when the shared variance is partitioned (Lahey et al. 2021; Moore and Lahey 2022). In the present study, we used the same bifactor specification and examined the general, specific internalizing, and specific externalizing factor scores; higher scores indicate greater mental health symptoms.

### 2.6 Additional Covariates and Sociodemographics

Additional covariates and sociodemographic variables included participants’ age (in months), caregiver-reported sex at birth, race/ethnicity, annual household income, highest parental education, perceived neighborhood safety, and physical activity. Caregivers reported participants’ race and ethnicity using a single item for race and a separate item for Hispanic/Latino ethnicity. Participants were classified into one of the following categories: White (non-Hispanic), Black (non-Hispanic), Hispanic/Latino, Asian/Pacific Islander, Multiracial (Non-Black), Multiracial (Black), and Other (non-Hispanic). Annual household income (“<$50,000 USD”, “≥ $50,000 - <$100,000 USD”, “≥$100,000 USD”, “Decline to answer”, or “Don’t know”) and highest parental education (“Up to high school”, “High school diploma/GED”, “Some college”, “Bachelor’s degree”, or “Graduate school or professional degree”) were each included as separate covariates using the listed categorical levels. Perceived neighborhood safety was assessed using a 3-item caregiver-reported neighborhood safety/crime survey from the PhenX Neighborhood Safety Protocol, evaluated on a 5-point Likert scale, where higher summary scores represented greater perceived neighborhood safety (Echeverria, Diez-Roux, and Link 2004; Mujahid et al. 2007). We included race/ethnicity, income, parental education, and perceived neighborhood safety to account for the higher exposure to ELA and air pollution often found among structurally disadvantaged groups (Hajat, Hsia, and O’Neill 2015; Nunez et al. 2024; Orendain et al. 2023). Finally, physical activity was assessed via the Youth Risk Behavior Survey and represented the number of days youth reported being physically active for a total of 60 minutes (Dolsen, Deardorff, and Harvey 2019; Hunsberger et al. 2015). Physical activity was included as a proxy for outdoor activity, which may be related to greater ambient air pollution exposure (Wu et al. 2024).

### 2.7 Statistical Analyses

Multiple linear mixed-effects (LMEs) analyses were conducted using R software (Version 4.5.1) (R Core Team 2025) to test the relationships among environmental risk factors (ELA and PM_2.5_), amygdala volumes (total and BLA subregion), and psychopathology symptoms (general, specific internalizing, specific externalizing factor scores). Covariates included: age, sex assigned at birth, highest caregiver education, annual household income, race/ethnicity, perceived neighborhood safety, self-reported physical activity, co-exposure to NO_2_ and O_3_, as well as total intracranial volume and handedness (the latter explicitly for analyses including the total amygdala and amygdala subregions). Each model included a random intercept for study site to account for site variability. First, we examined variance inflation factors (VIFs) to assess multicollinearity among predictors; all VIFs were < 2 (**Supplemental Table 2**). For our first aim, total amygdala and each BLA subregion volume (LA, BLDI, BM, BLVPL) were modeled as independent outcomes across five LMEs. Each of these models assessed the main effects of ELA and PM_2.5_ total mass and their interaction, to evaluate whether PM_2.5_ total mass moderated the association between ELA and amygdala volumes. In our second aim, using the same approach, general psychopathology, specific internalizing, and specific externalizing symptoms were each modeled as separate outcomes across three LMEs. Each of these models included main effects of ELA, PM_2.5_ total mass, and their interaction, to evaluate whether PM_2.5_ total mass moderated the association between ELA and psychopathology symptoms. Lastly, to test whether amygdala volumes, ELA, and PM_2.5_ total mass jointly relate to psychopathology, each psychopathology factor score (general, specific internalizing, or specific externalizing) was modeled as three separate outcomes. These models included main effects of amygdala volumes, ELA, and PM_2.5_ total mass, along with their two-way (Amygdala × ELA, Amygdala × PM_2.5_, ELA × PM_2.5_) and three-way (Amygdala × ELA × PM_2.5_) interaction terms, which yielded 15 total models. Using a model-building approach, the full model including the three-way interaction was tested; if the three-way interaction was not significant and did not improve model fit, the more parsimonious two-way interaction model was retained. Model fit was compared using likelihood-ratio tests, Akaike information criterion (AICs), and Bayesian information criterion (BICs).

Post-hoc analyses were implemented to characterize significant interactions. Specifically, for interactions between ELA and PM_2.5_, the simple slopes were estimated at the mean of PM_2.5_ (7.52 μg/m³), and one standard deviation above and below the mean (9.00 μg/m³ and 6.04 μg/m³). For interactions between amygdala and ELA, simple slopes were estimated at no ELA exposure (0 counts), mean ELA exposure (1.43 counts), and one standard deviation above the mean (2.87 counts). False discovery rate (FDR) corrections were applied across outcomes within each set of analyses (Aim 1: five amygdala volumes and Aims 2 and 3: three p-factor outcomes). Analyses were conducted using mean-centered PM_2.5_ and z-scored amygdala brain volumes to help facilitate interpretation across regions.

To complement our LME analyses, we also conducted exploratory weighted quantile sum (WQS) regression analyses as previously published (Sukumaran et al. 2024) in order to disentangle whether a mixture of the 15 PM_2.5_ components (Br, Cu, Fe, EC, OC, NH₄⁺, NO_3_⁻, SO₄^2^⁻, V, Zn, Ni, Pb, Ca, Si, K) moderated the relationship between ELA and psychopathology symptoms and amygdala volumes. WQS regression is a mixture analysis method implemented within a generalized linear regression framework that combines quantile mixture components into a single weighted index (WQS index), with data-derived weights showing the relative importance of each mixture component to the WQS index. The 15 PM_2.5_ components were scored into deciles within the WQS analysis conducted using the gWQS package (Renzetti et al. 2021) with models fit using the gwqs() function, using 500 bootstrap samples and a 60% site-stratified validation split. WQS requires specification of a direction for associations between the overall mixture and the outcome as well as a direction for the interaction effect for models with interaction terms. We had no *a priori* directionality assumption in how the weighted mixture and ELA would interact to relate to our outcomes of interest. Thus, each outcome was fit with WQS (15 PM_2.5_ components) × ELA interaction models for each main-effect direction, evaluating both positive and negative interactions. We included the same covariates as LME models and included site as a fixed effect. FDR correction was done within each outcome group (amygdala or psychopathology symptom scores).

## 3. Results

The full analytic sample and details are presented in **Table 1**, and comparisons to the larger ABCD Study sample at baseline are presented in **Supplemental Table 3**. Broadly, compared to the ABCD Study cohort at baseline, our analyzed subsample was more likely to be non-Hispanic White participants from higher-income and more educated households than other ABCD participants. For the exposures of interest, average PM_2.5_ was modestly lower in the analytic sample, whereas ELA was comparable across the two samples. Psychopathology bifactor scores and amygdala volumes were nearly identical between the two samples.

**Table 1.**
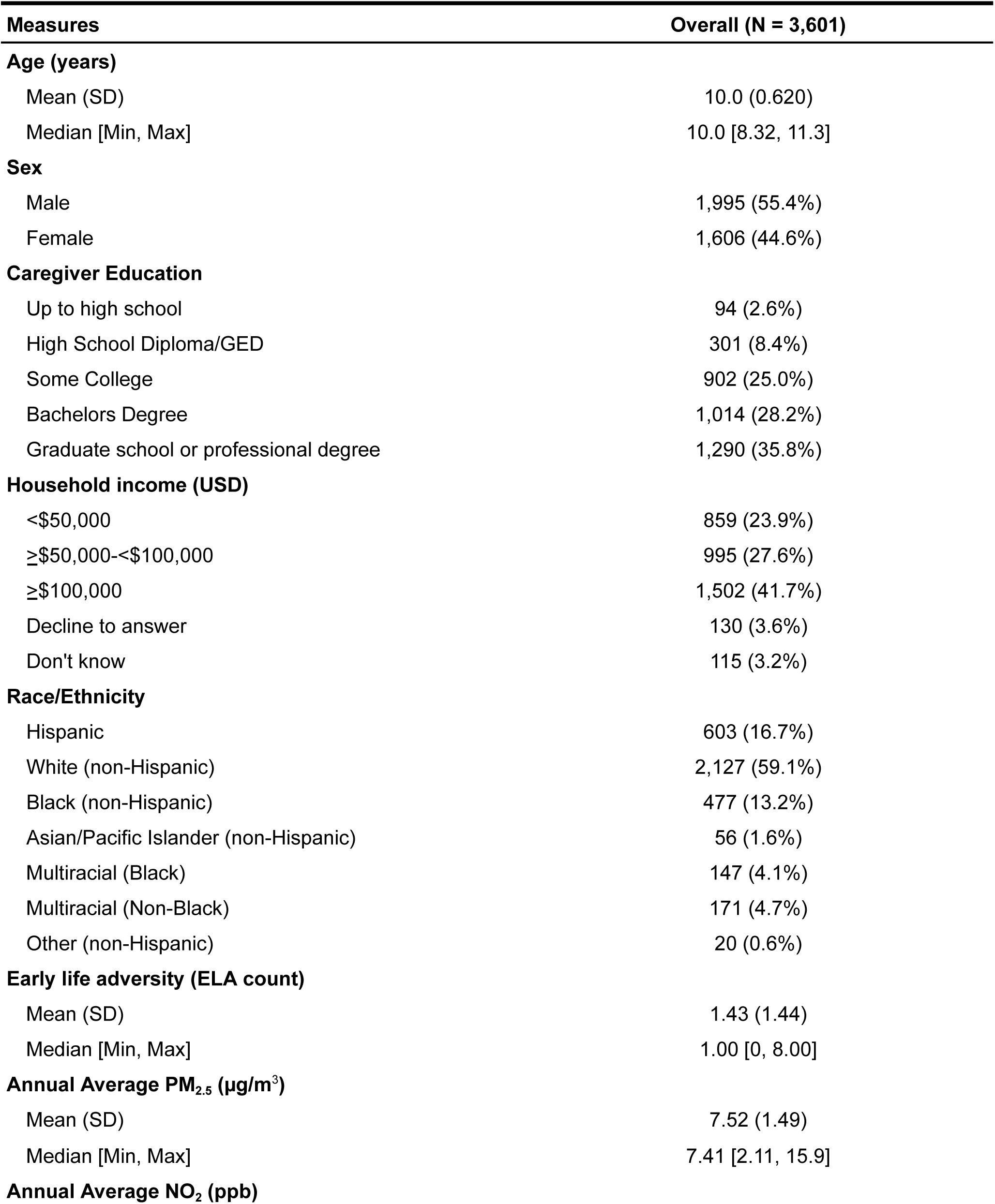
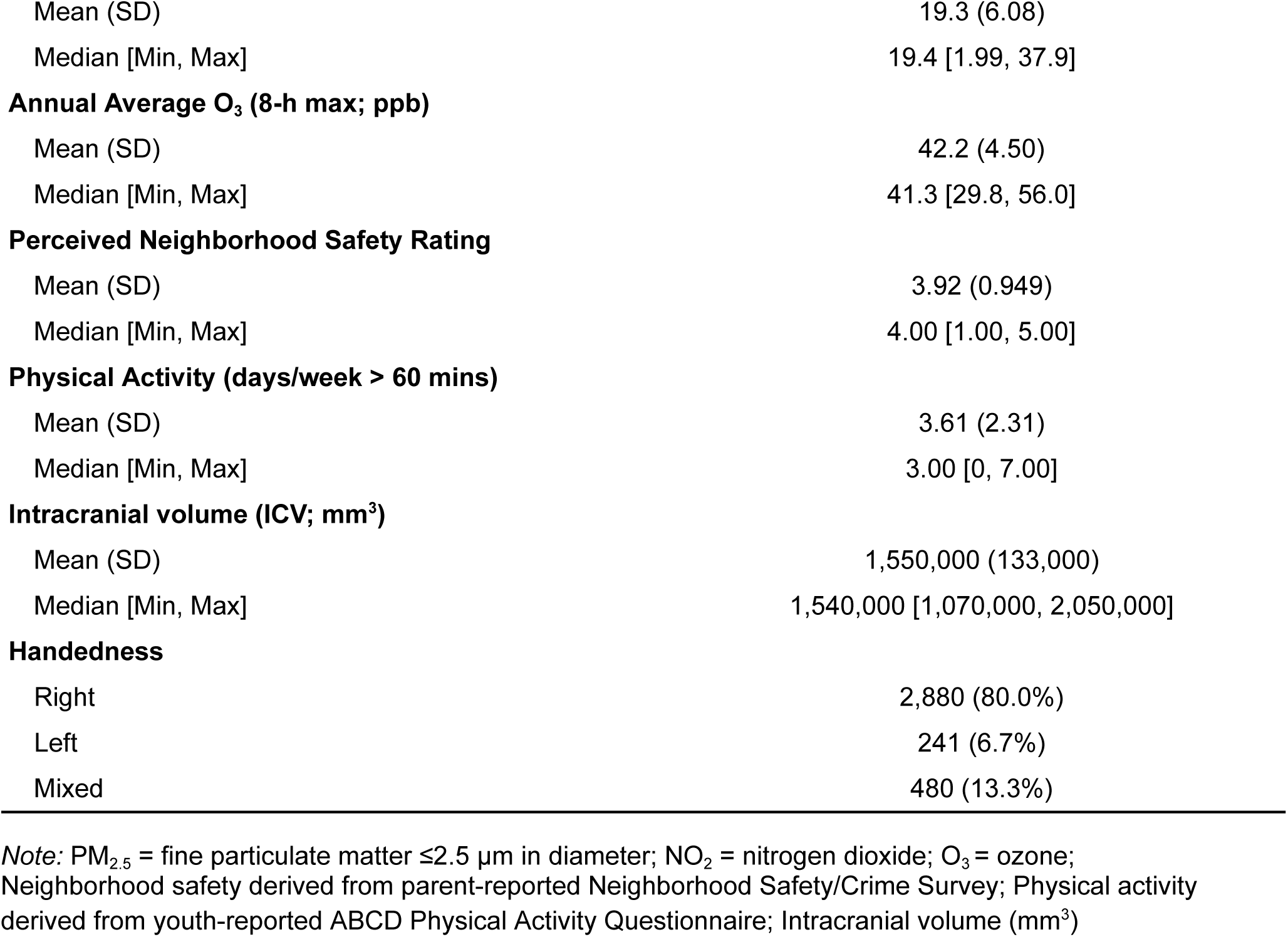
Demographic, Environmental Exposures, and Brain Characteristics of the Analytic Sample.

| Measures | Overall (N = 3,601) |
| --- | --- |
| **Age (years)** |  |
| Mean (SD) | 10.0 (0.620) |
| Median [Min, Max] | 10.0 [8.32, 11.3] |
| **Sex** |  |
| Male | 1,995 (55.4%) |
| Female | 1,606 (44.6%) |
| **Caregiver Education** |  |
| Up to high school | 94 (2.6%) |
| High School Diploma/GED | 301 (8.4%) |
| Some College | 902 (25.0%) |
| Bachelors Degree | 1,014 (28.2%) |
| Graduate school or professional degree | 1,290 (35.8%) |
| **Household income (USD)** |  |
| <\$50,000 | 859 (23.9%) |
| ≥\$50,000-<\$100,000 | 995 (27.6%) |
| ≥\$100,000 | 1,502 (41.7%) |
| Decline to answer | 130 (3.6%) |
| Don't know | 115 (3.2%) |
| **Race/Ethnicity** |  |
| Hispanic | 603 (16.7%) |
| White (non-Hispanic) | 2,127 (59.1%) |
| Black (non-Hispanic) | 477 (13.2%) |
| Asian/Pacific Islander (non-Hispanic) | 56 (1.6%) |
| Multiracial (Black) | 147 (4.1%) |
| Multiracial (Non-Black) | 171 (4.7%) |
| Other (non-Hispanic) | 20 (0.6%) |
| **Early life adversity (ELA count)** |  |
| Mean (SD) | 1.43 (1.44) |
| Median [Min, Max] | 1.00 [0, 8.00] |
| **Annual Average PM<sub>2.5</sub> (µg/m<sup>3</sup>)** |  |
| Mean (SD) | 7.52 (1.49) |
| Median [Min, Max] | 7.41 [2.11, 15.9] |
| **Annual Average NO<sub>2</sub> (ppb)** |  |
| Mean (SD) | 19.3 (6.08) |
| Median [Min, Max] | 19.4 [1.99, 37.9] |
| **Annual Average O<sub>3</sub> (8-h max; ppb)** |  |
| Mean (SD) | 42.2 (4.50) |
| Median [Min, Max] | 41.3 [29.8, 56.0] |
| **Perceived Neighborhood Safety Rating** |  |
| Mean (SD) | 3.92 (0.949) |
| Median [Min, Max] | 4.00 [1.00, 5.00] |
| **Physical Activity (days/week > 60 mins)** |  |
| Mean (SD) | 3.61 (2.31) |
| Median [Min, Max] | 3.00 [0, 7.00] |
| **Intracranial volume (ICV; mm<sup>3</sup>)** |  |
| Mean (SD) | 1,550,000 (133,000) |
| Median [Min, Max] | 1,540,000 [1,070,000, 2,050,000] |
| **Handedness** |  |
| Right | 2,880 (80.0%) |
| Left | 241 (6.7%) |
| Mixed | 480 (13.3%) |

*Note:* PM<sub>2.5</sub> = fine particulate matter $\leq 2.5$ $\mu\text{m}$ in diameter; NO<sub>2</sub> = nitrogen dioxide; O<sub>3</sub> = ozone; Neighborhood safety derived from parent-reported Neighborhood Safety/Crime Survey; Physical activity derived from youth-reported ABCD Physical Activity Questionnaire; Intracranial volume (mm<sup>3</sup>)

Our sample participants (N = 3,601) were approximately 10 years old (SD=0.62) and reported experiencing a mean of 1.42 ELAs (SD=1.44, range= 0-8). Annual average PM_2.5_ exposure levels for the current sample were 7.49 µg/m^3^ (SD=1.49), which was significantly lower than current 2024 Environmental Protection Agency (EPA) standards (i.e., 9 µg/m^3^; t(3600) = -60.81, p < 0.001), but significantly higher than the World Health Organization (WHO) 2021 Air Quality Guidelines (i.e., 5 µg/m^3^; (t(3600) = 100.28, p < 0.001).

### 3.1 ELA, PM_2.5_, and Amygdala Volumes

Using linear mixed-effects modeling, neither the ELA × PM_2.5_ interaction nor the main effects of ELA or PM_2.5_ were significant for either the total amygdala volume or any of the BLA subregions following FDR correction (**Supplemental Table 4**). Exploratory mixture models for the amygdala volumes also yielded no significant ELA × WQS interactions or main effects (**Supplemental Table 5**).

### 3.2 ELA, PM_2.5_, and Psychopathology Symptoms

The ELA × PM_2.5_ interaction was significant for the specific externalizing factor (*β* = 0.005, 95% *CI* [0.002-0.008], FDR-*p* = 0.001; **Supplemental Table 6**), but not for general psychopathology or the specific internalizing factor. Post-hoc simple slope analyses indicated that the positive association between ELA and the specific externalizing factor increased in magnitude at higher levels of annual PM_2.5_ exposure (**Figure 2, Supplemental Table 7**). Additionally, ELA had significant main effects for all three psychopathology symptom outcomes (**Supplemental Table 6**). Together, these findings suggest that ELA is associated with psychopathology symptoms and the association between ELA and externalizing was *stronger* among children with *higher* PM_2.5_ total mass exposure.

**Fig. 2.**
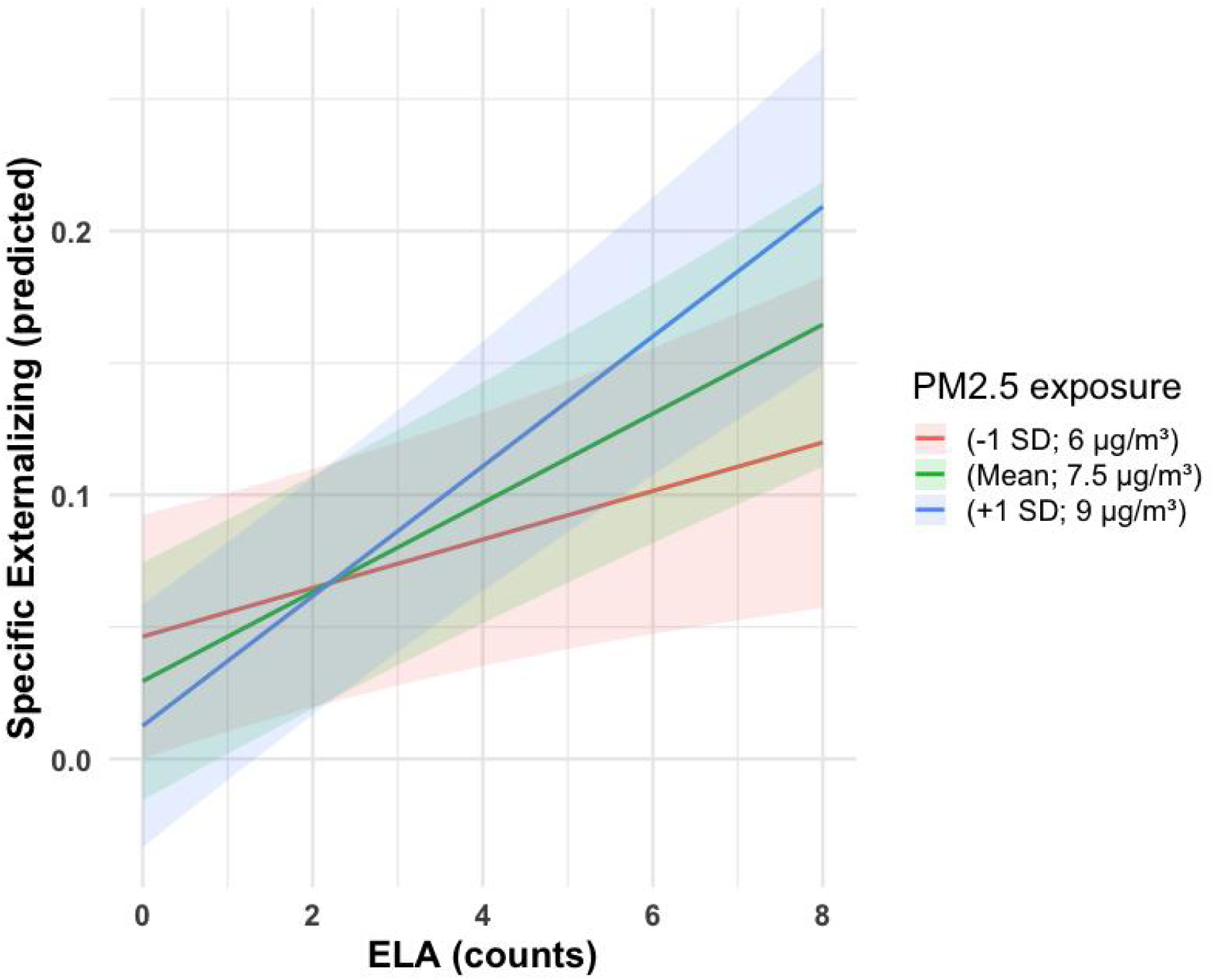
Significant interaction between ELA and PM_2.5_ on externalizing symptoms. PM_2.5_ moderated the association between ELA and externalizing, such that youth with higher PM_2.5_ exposure and greater ELA counts had greater specific externalizing symptoms (illustrated by blue line).

In our exploratory WQS models, examining co-exposure to the 15 PM_2.5_ components as a moderator of ELA, we found no significant ELA by WQS interactions, nor main effect of the PM_2.5_ component mixture, for any psychopathology factor scores (**Supplemental Table 8**).

### 3.3 Interactions between ELA, PM_2.5_, and Amygdala on Psychopathology Symptoms

To examine how amygdala volumes, ELA, and PM_2.5_ interact to relate to psychopathology, we investigated all interactions within the same model. Per our model-building approach, the three-way interaction (Amygdala × ELA × PM_2.5_) was not significant, and the two-way interaction models were more parsimonious (14/15 models, **Supplemental Table 9**). There was a significant BLVPL volume by ELA interaction for general psychopathology (**Figure 3A**; *β* = -0.013, 95% *CI* [-0.023-0.002], FDR-*p* = 0.03; **Supplemental Table 10**) and specific externalizing (**Figure 3B**; *β* = -0.006, 95% *CI* [-0.01 - -0.001], FDR-*p* = 0.03; **Supplemental Table 11**), but not for specific internalizing symptoms (**Supplemental Table 12**). Specifically, post-hoc simple slope analyses for general psychopathology symptoms indicated that *smaller* BLVPL volume was associated with *greater* general psychopathology among youth with *higher* ELA exposure, whereas this association was weaker at the mean ELA exposure and absent among youth with no ELA exposure (**Supplemental Table 13**). For specific externalizing, post-hoc simple slopes showed a similar directional effect: *smaller* BLVPL volume was associated with *greater* externalizing symptoms at *higher* ELA exposure (**Supplemental Table 14**). Other significant results were consistent with the patterns reported above, including main effects of ELA on psychopathology symptoms in models including the total amygdala and other BLA volumes (**Supplemental Tables 10-12**) as well as the PM_2.5_ x ELA interaction on externalizing symptoms across all models including amygdala volumes (**Supplemental Table 11**).

**Fig. 3.**
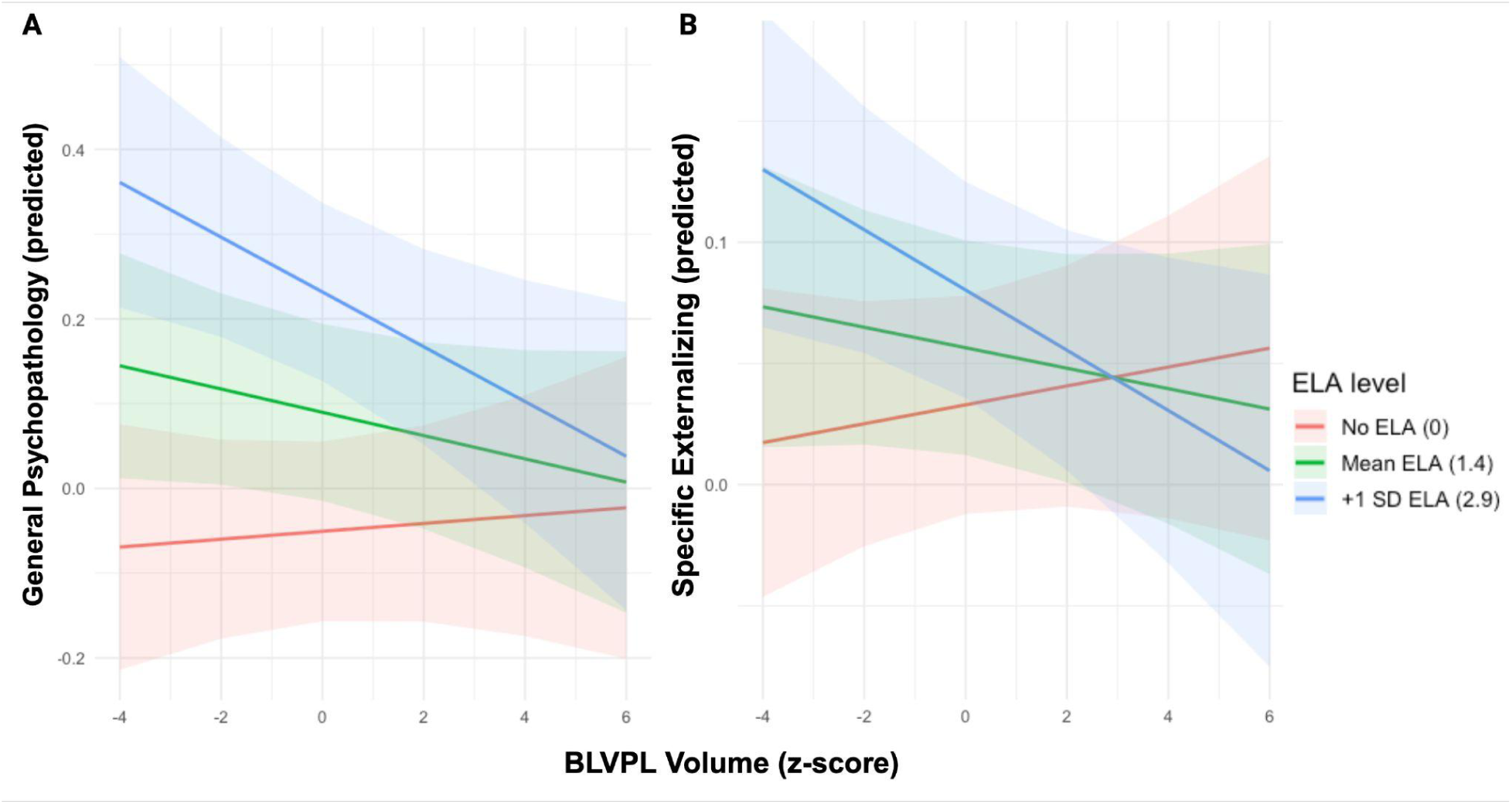
Significant interaction between ELA and BLVPL on A) general psychopathology symptoms and B) specific externalizing symptoms. ELA moderated the association between BLVPL and generalized psychopathology and externalizing symptoms, such that youth with higher ELA exposure (shown by blue line, +1 SD above the mean of ELA) showed a stronger negative association between BLVPL volume and mental health symptoms.

## 4. Discussion

The present study examined three independent moderation models examining the unique and interactive associations of environmental exposures (ELA and annual average PM_2.5_ exposure) with amygdala structure and psychopathology in 9- to 10-year-old children. Consistent with prior work and our hypotheses, ELA was associated with greater symptoms across all psychopathology domains, including general, specific externalizing, and specific internalizing symptoms (Wade et al. 2022). Supporting our hypotheses, PM_2.5_ exposure moderated the association between ELA and specific externalizing symptoms, such that youth with both higher ELA and higher PM_2.5_ exposure exhibited the greatest specific externalizing scores at ages 9-10 years. These findings are aligned with the ‘double hit’ framework (Clougherty and Kubzansky 2009), suggesting combined physical and psychosocial environmental risk factors may be associated with greater mental health problems. Against our hypotheses, we did not observe a direct negative effect of ELA on BLA volumes, nor was the association moderated by PM_2.5_. However, we found that smaller BLVPL volumes were associated with greater general and specific externalizing scores among youth exposed to higher ELAs. In exploratory mixture analyses, none of the 15 PM_2.5_ components showed significant main effects or interactions. This suggests that associations with PM_2.5_ total mass in our primary analyses may be driven by total particle matter (regardless of their composition), and by constituents not captured among the 15 measured components, or both. Together, these findings suggest that ELA is related to higher psychopathology symptoms in preadolescence, co-occurring PM_2.5_ total mass exposure may compound the ELA-related risk for externalizing symptoms, and smaller BLVPL volumes were associated with heightened vulnerability to ELA’s effects on general and specific externalizing factor scores.

Consistent with a robust body of prior work, we found that greater ELA exposure was associated with greater psychopathology symptoms. This main effect of ELA remained significant across all models, even as ELA interacted with PM_2.5_ exposure and BLVPL volume to further shape individual differences in preadolescent specific externalizing and general psychopathology symptoms. While we found ELA to interact with PM_2.5_ and increase risk for externalizing symptoms, ELA also appears to operate *independently* in its association with internalizing and general psychopathology symptomology at this age of development. This is broadly consistent with recent population neuroscience efforts that incorporate multi-dimensional exposure data into large predictive models of psychopathology risk, many of which find that proximal, day-to-day experiences carry greater predictive weight compared to more distal environmental exposures or neurobiological markers (Moore et al. 2022; Pagliaccio et al. 2024). Our findings underscore that understanding mental health risk requires integrating psychosocial and physical risk factors, rather than considering ambient environmental exposures in isolation, as these risk factors may operate along independent pathways that converge on shared outcomes.

Beyond the main effect association of ELA, we found that PM_2.5_ exposure moderated the relationship between ELA and externalizing scores, such that youth with both higher ELA and higher PM_2.5_ exposure exhibited the greatest specific externalizing scores. This pattern is consistent with a ‘double hit’ framework of environmental risk, in which early psychosocial adversity sensitizes the developing stress system such that subsequent physical environmental exposures here, ambient air pollution, confer disproportionately greater risk than either exposure alone. This finding aligns with prior work showing that combined psychosocial and physical environmental stressors may magnify risk for psychopathology beyond either exposure individually (Pagliaccio et al. 2020; Vargas et al. 2025; Zundel et al. 2022). Our study extends this literature by using a transdiagnostic bifactor model to separate general psychopathology risk from symptom-specific variance. We found that the ‘double hit’ pattern was unique to externalizing-specific symptoms in a large, sociodemographically diverse sample of 9- to 10-year-olds. This interaction did not extend to specific internalizing or general psychopathology, suggesting that the combined risk of ELA and PM_2.5_ exposure may be especially relevant to externalizing-specific symptoms, including inattention, impulsivity, and behavioral and emotional dysregulation. Notably, when examined independently of psychosocial adversity, PM_2.5_ exposure has shown inconsistent, and at times counterintuitive, associations with behavioral problems in this same cohort, including a longitudinal study finding that higher PM_2.5_ and NO_2_ exposure at ages 9-10 was associated with negligible behavioral problems over a subsequent 2-year follow-up (Campbell et al. 2024) and another showing only the number of days of higher exposure in a year was associated with increases in internalizing problems in the ABCD cohort (Smolker et al. 2024). Moreover, PM_2.5_ is a heterogeneous mixture whose composition depends on source (Dominici et al. 2015; Sangkham 2026). Our mixture analyses largely corroborated our primary LME findings. PM_2.5_ component mixture was not independently significantly associated with general psychopathology, specific internalizing, or specific externalizing. This contrasts with our primary PM_2.5_ total mass models, where PM_2.5_ total mass moderated ELA’s relationship with the specific externalizing factor. This suggests that for the variance unique to externalizing symptoms, the total aggregate dose of fine particles, rather than the specific composition captured by these 15 components, may be what matters for mental health risk. Because PM_2.5_ total mass integrates constituents beyond those 15 measured components (Dominici et al. 2015), a signal residing in this unmeasured fraction would surface in the total mass models but not in the component mixture models. Importantly, our study sample also represented only a fraction of the larger ABCD cohort, because the CIT168 atlas was developed using Siemens-acquired data, and its performance on non-Siemens data is unknown. Because of this exclusion of non-Siemens study sites, our sample also differed from the broader ABCD cohort on several PM_2.5_ components, including NH_4_ ^+^, NO_3_ ^-^, SO ^2^, and V (**Supplemental Table 3**), suggesting that our mixture analyses may not have captured the fuller range of PM_2.5_ mixtures observed more broadly across the U.S. Together, with the present findings, this suggests that PM_2.5_’s association with externalizing behavior may not be reliably detectable as a main effect in this relatively low-exposure cohort, but instead may emerge specifically in the context of co-occurring psychosocial adversity, underscoring the value of modeling these exposures jointly rather than in isolation.

Contrary to our hypotheses, ELA did not show a direct relationship with amygdala volumes. This is not entirely surprising given that prior work on ELA, PM_2.5_, and amygdala volume has been inconsistent, with studies reporting positive, negative, and null findings (Hanson and Nacewicz 2021; McLaughlin et al. 2019). This inconsistency may partly reflect the specific type of ELA examined, as dimensional models of adversity suggest that threat- and deprivation-related ELA differentially impact brain regions, systems, and speed of maturation (e.g., accelerated versus delayed) (McLaughlin et al. 2019). The present study examined cumulative ELA counts rather than distinguishing between threat and deprivation subtypes, which may have obscured subtype-specific associations with amygdala structure. Although this study was conducted in children with a relatively narrow age range, this discrepancy may also, in part, reflect the unique, protracted developmental trajectory of the amygdala throughout childhood and into early adulthood (Russell, Marsee, and Weems 2021; Wierenga et al. 2014). Prior work suggests that associations between ELA and amygdala volume may emerge at different developmental stages, and we may have been unable to capture these specific associations in our baseline sample. For example, lower family income and limited parental education were associated with smaller amygdala volume in middle and late adolescents (13–21 years) but not younger children (3–12 years) (Merz, Tottenham, and Noble 2018), while greater threat exposure (e.g., physical, emotional, and sexual abuse, domestic violence) was associated with smaller amygdala volumes specifically among younger (8-11 years), but not older, participants (15-17 years) (Peverill et al. 2023). Sample severity may also play a role in our null findings being that many prior studies documenting amygdala differences sampled youth with more severe ELA exposure (>3 ELAs), whereas ABCD is a community sample in which more than half of our baseline sample reported minimal ELA exposure. Together, these factors may help explain inconsistency in ELA-amygdala volume associations. Future work should aim to capture the evolving, longitudinal relationship between ELA and brain development, while also accounting for distinct types of ELA exposure.

Partially in line with our hypotheses, we found smaller BLVPL volumes were associated with greater psychopathology symptoms (general and externalizing) among youth exposed to *higher* ELAs. The specificity of the relationship between the BLVPL and psychopathology, rather than any other amygdala subregion, may reflect its unique developmental profile relative to its counterparts. The BLVPL encompasses the paralaminar nucleus (deCampo and Fudge 2012), which is defined by its continued maturation well beyond childhood (Ojha et al. 2025; Sorrells et al. 2018), and contains immature, migratory excitatory neurons that persist into adulthood, with transcriptional and structural changes emerging specifically during adolescence (Avino et al. 2018; Page et al. 2022; Sorrells et al. 2018). This protracted developmental trajectory has led some to believe that the paralaminar nucleus functions as a reservoir of ongoing amygdala-hippocampal neuroplasticity across adolescence (Sorrells et al. 2018). Our previous work in the baseline ABCD study cohort has similarly highlighted the BLVPL as a subregion of importance for development, showing robust sex-specific patterns of volume and apportionment not observed in other amygdala subregions (Overholtzer et al. 2025). Given the BLVPL’s extended developmental window, it may be especially sensitive to the influence of early psychosocial experience, which could help explain why its association with psychopathology, but not that of other amygdala subregions, depended on level of ELA exposure. The specificity of our effect to the variance shared by all symptoms (operationalized by the general factor) and the remaining variance specific to externalizing symptoms may partly reflect measurement factors. Externalizing behaviors are more behaviorally observable in the baseline sample (9-10-year-olds). Thus, externalizing may be more reliably captured by caregiver reports than internalizing, which may be underreported or under-detected by caregivers (our informants) (Baumgartner et al. 2020; Carvalho, Koss, and Ravindran 2026). Additionally, our findings may reflect typical developmental differences: ELA’s associations with externalizing/behavioral dysregulation tend to emerge earlier in childhood, while associations with internalizing symptoms often increase later in adolescence. Specifically, internalizing symptoms typically show a marked increase in prevalence and severity beginning around puberty onset (Alloy et al. 2016; Pfeifer and Allen 2021). Given the majority of children in this ABCD baseline and Siemens-only subsample were at pre- or early-pubertal stages (Overholtzer et al. 2025), internalizing symptoms may not yet have been sufficiently expressed at this developmental stage to reveal associations, which may emerge later in adolescence. Future work incorporating longitudinal data may help untangle whether ELA and BLVPL associations exist for internalizing symptoms later in adolescence.

The present study leverages several key strengths, including a large, sociodemographically heterogeneous sample from the ABCD Study, the use of the CIT168 probabilistic atlas to characterize functionally and developmentally distinct amygdala subregions rather than whole-structure volume alone, integrating psychosocial and physical environmental exposures, and a bifactor model of psychopathology that captures both transdiagnostic (general) and domain-specific (internalizing, externalizing) symptom variance. Several limitations should also be noted. First, the present study is cross-sectional, which precludes conclusions about temporal precedence or causality among ELA, PM_2.5_ exposure, amygdala structure, and psychopathology. For example, while we interpret smaller BLVPL volume as a substrate through which ELA confers risk for psychopathology symptoms, it is equally plausible that early psychopathology symptoms influence subsequent brain development, or that a third, unmeasured factor drives both. Thus, this lack of temporal ordering in our exposures and outcomes limit our ability to formally test a moderated mediation model. Second, we operationalized ELA as a cumulative count of adverse experiences, which does not distinguish between types of adversity, though different adversity types have been shown to differentially affect brain structure. Future work incorporating both frequency and dimensional models of ELA may better capture these subtype-specific pathways. Third, we focused on specific amygdala subregions *a priori* given their theorized sensitivity to early environmental input, but ELA has been linked to structural and functional differences across broader corticolimbic and frontoparietal networks. Future studies should examine whether the patterns observed here extend to other neural systems implicated in adversity-related risk. Finally, the amygdala segmentation process was normalized on Siemens scanners only; thus our analytic sample was higher-SES, disproportionately White, and regionally limited (**Supplemental Table 3**). In contrast, a recent study investigating air pollution exposure with a less restrictive ABCD sample uncovered air pollution differences by race/ethnicity and income vary substantially by geographic site (Cardenas-Iniguez et al. 2025), suggesting that the reported findings here may underestimate those that would emerge in more representative samples. Future longitudinal research, leveraging the ongoing follow-up waves of the ABCD Study, is needed to clarify the direction and developmental timing of these associations, including whether ELA-related alterations in BLVPL structure precede the onset of psychopathology symptoms, whether these associations strengthen or attenuate across adolescence as the BLVPL continues to mature, and whether the ‘double-hit’ pattern observed for PM_2.5_ and ELA on externalizing symptoms persists, emerges, or resolves over time and if these findings are similar for internalizing at later ages. Incorporating repeated measures of ELA, air pollution exposure, amygdala structure, and psychopathology along with dimensional ELA measures and broader neural network involvement, across development would allow for within-person modeling of change and a more comprehensive test of these proposed mechanisms.

## 5. Conclusions

We provide evidence that psychosocial and physical environmental factors are jointly related to psychopathology symptoms in preadolescence. ELA was robustly associated with greater psychopathology symptoms, while co-occurrence with PM_2.5_ exposure specifically moderated the ELA relationship on the specific externalizing factor of symptoms. Additionally, smaller BLVPL volume was related to greater general and specific externalizing symptoms among youth exposed to higher ELA. Together, while previous research showed that ELA and PM_2.5_ exposures independently relate to increased mental health risk, our findings suggest that ELA and PM_2.5_ may also interact and amplify mental health risk during preadolescence. As this study is cross-sectional, we cannot establish the direction of these associations; thus, longitudinal work is needed to evaluate these relationships across development. Broadly, these results underscore the value of integrating psychosocial, physical, and neurobiological factors when identifying children at risk for psychopathology.

## Acknowledgments

A special thank you to all participants and their families for their participation in the ABCD Study. We would also like to acknowledge Jade Li, who completed preliminary descriptives and simple analyses between the amygdala and p-factor scores for her undergraduate thesis project in 2025.

## Pre-registration statement

This study was not pre-registered.

## Funding statement

Research described in this article was supported by the National Institutes of Health grants [NIEHS: R01ES032295, R01ES031074, P30ES007048-23S1, 3P30ES000002-55S1, U24ES036819; NIA: T32AG000037; NIMH: F30MH139336].

Data used in the preparation of this article were obtained from the Adolescent Brain Cognitive Development® (ABCD) Study (https://abcdstudy.org), held in the NIMH Data Archive (NDA). This is a multisite, longitudinal study designed to recruit more than 10,000 children aged 9-10 and follow them over 10 years into early adulthood. The ABCD Study® is supported by the National Institutes of Health and additional federal partners under award numbers U01DA041048, U01DA050989, U01DA051016, U01DA041022, U01DA051018, U01DA051037, U01DA050987, U01DA041174, U01DA041106, U01DA041117, U01DA041028, U01DA041134, U01DA050988, U01DA051039, U01DA041156, U01DA041025, U01DA041120, U01DA051038, U01DA041148, U01DA041093, U01DA041089, U24DA041123, U24DA041147. A full list of supporters is available at https://abcdstudy.org/federal-partners.html. A listing of participating sites and a complete listing of the study investigators can be found at https://abcdstudy.org/consortium_members/. ABCD consortium investigators designed and implemented the study and/or provided data but did not necessarily participate in the analysis or writing of this report. This manuscript reflects the views of the authors and may not reflect the opinions or views of the NIH or ABCD consortium investigators. Additional support for this work was made possible from NIEHS R01-ES032295 and R01-ES031074.

The ABCD data repository grows and changes over time. The ABCD data used in this report came from the ABCD 3.0, 5.1 and 6.1 data releases. DOIs can be found at ABCD Release 3.0; https://nda.nih.gov/study.html?id=901; ABCD Release 5.1; https://nda.nih.gov/study.html?id=2313; and ABCD Release 6.1; https://www.nbdc-datahub.org/abcd-release-6-1.

## Data availability

The authors do not have permission to share the data. Due to the sensitive nature of this study and in accordance with ABCD Study guidelines, raw data from this study will not be made available. The dataset used in this study can be obtained with authorization from https://nda.nih.gov/abcd and at https://www.nbdc-datahub.org/abcd-release-6-1. Application and approval of an NDA form are required.

## CReDIT authorship contributions

**Amanda C. Del Giacco**: Conceptualization, Data curation, Methodology, Formal analysis, Project administration, Writing - Original draft & editing, Visualization. **Michael R. Rosario**: Conceptualization, Data curation, Methodology, Formal analysis, Project administration, Writing - Original draft & editing, Visualization. **Carlos Cardenas-Iniguez**: Methodology, Data curation, Writing - Review & editing. **Nitya Chawla**: Visualization, Writing - Review & editing. **Alan Wen**: Visualization, Writing - Review & editing**. Kirthana Sukamaran**: Methodology, Data curation, Writing - Review & editing. **L. Nate Overholtzer**: Data curation, Writing - Review & editing. **Jiu-Chiuan Chen:** Funding acquisition, Methodology, Writing - Review & editing. **Benjamin B. Lahey**: Methodology, Software, Writing - Review & editing. **Tyler M. Moore**: Methodology, Software, Writing - Review & editing. **Megan M. Herting**: Funding acquisition, Conceptualization, Methodology, Supervision, Project Administration, Writing - Original & editing.

## References

Achenbach, Thomas M., and LA Rescorla. 2001. Child Behavior Checklist for Ages 6-18. University of Vermont Burlington, VT.

Ahmed, Salma M., Gita D. Mishra, Katrina M. Moss, Ian A. Yang, Kate Lycett, and Luke D. Knibbs. 2022. “Maternal and Childhood Ambient Air Pollution Exposure and Mental Health Symptoms and Psychomotor Development in Children: An Australian Population-Based Longitudinal Study.” Environment International 158:107003. doi:10.1016/j.envint.2021 **Alan Wen:** Visualization, Writing - Review & editing..107003.

Alloy, Lauren B., Jessica L. Hamilton, Elissa J. Hamlat, and Lyn Y. Abramson. 2016. “Pubertal Development, Emotion Regulatory Styles, and the Emergence of Sex Differences in Internalizing Disorders and Symptoms in Adolescence.” Clinical Psychological Science 4(5):867–81. doi:10.1177/2167702616643008.

Avino, Thomas A., Nicole Barger, Martha V. Vargas, Erin L. Carlson, David G. Amaral, Melissa D. Bauman, and Cynthia M. Schumann. 2018. “Neuron Numbers Increase in the Human Amygdala from Birth to Adulthood, but Not in Autism.” Proceedings of the National Academy of Sciences 115(14):3710–15. doi:10.1073/pnas.1801912115.

Azad, Anisa, Ryan P. Cabeen, Farshid Sepehrband, Robert Kim, Claire E. Campbell, Kirsten Lynch, J. Michael Tyszka, and Megan M. Herting. 2021. “Microstructural Properties within the Amygdala and Affiliated White Matter Tracts across Adolescence.” NeuroImage 243:118489. doi:10.1016/j.neuroimage.2021.118489.

Baumgartner, Noemi, Isabelle Häberling, Sophie Emery, Michael Strumberger, Kristin Nalani, Suzanne Erb, Silke Bachmann, Lars Wöckel, Ulrich Müller-Knapp, Bruno Rhiner, Brigitte Contin-Waldvogel, Klaus Schmeck, Susanne Walitza, and Gregor Berger. 2020. “When Parents and Children Disagree: Informant Discrepancies in Reports of Depressive Symptoms in Clinical Interviews.” Journal of Affective Disorders 272:223–30. doi:10.1016/j.jad.2020.04.008.

Becker, Léa J., Clémentine Fillinger, Robin Waegaert, Sarah H. Journée, Pierre Hener, Beyza Ayazgok, Muris Humo, Meltem Karatas, Maxime Thouaye, Mithil Gaikwad, Laetitia Degiorgis, Marie des Neiges Santin, Mary Mondino, Michel Barrot, El Chérif Ibrahim, Gustavo Turecki, Raoul Belzeaux, Pierre Veinante, Laura A. Harsan, Sylvain Hugel, Pierre-Eric Lutz, and Ipek Yalcin. 2023. “The Basolateral Amygdala-Anterior Cingulate Pathway Contributes to Depression-like Behaviors and Comorbidity with Chronic Pain Behaviors in Male Mice.” Nature Communications 14(1):2198. doi:10.1038/s41467-023-37878-y.

Bradley, Melissa, Kimberlie Dean, Samsung Lim, Kristin R. Laurens, Felicity Harris, Stacy Tzoumakis, Kirstie O’Hare, Vaughan J. Carr, and Melissa J. Green. 2024. “Early Life Exposure to Air Pollution and Psychotic-like Experiences, Emotional Symptoms, and Conduct Problems in Middle Childhood.” Social Psychiatry and Psychiatric Epidemiology 59(1):87–98. doi:10.1007/s00127-023-02533-w.

Breslin, Florence J., Erin L. Ratliff, Zsofia P. Cohen, Julie M. Croff, and Kara L. Kerr. 2025. “Measuring Adversity in the ABCD® Study: Systematic Review and Recommendations for Best Practices.” BMC Medical Research Methodology 25(1):77. doi:10.1186/s12874-025-02521-5.

Brieant, Alexis, Anna Vannucci, Hajer Nakua, Jenny Harris, Jack Lovell, Divya Brundavanam, Nim Tottenham, and Dylan G. Gee. 2023. “Characterizing the Dimensional Structure of Early-Life Adversity in the Adolescent Brain Cognitive Development (ABCD) Study.” Developmental Cognitive Neuroscience 61:101256. doi:10.1016/j.dcn.2023.101256.

Bzdok, Danilo, Angela R. Laird, Karl Zilles, Peter T. Fox, and Simon B. Eickhoff. 2013. “An Investigation of the Structural, Connectional, and Functional Subspecialization in the Human Amygdala.” Human Brain Mapping 34(12):3247–66. doi:10.1002/hbm.22138.

Callaghan, Bridget L., and Nim Tottenham. 2016. “The Stress Acceleration Hypothesis: Effects of Early-Life Adversity on Emotion Circuits and Behavior.” Development and Behavior 7:76–81. doi:10.1016/j.cobeha.2015.11.018.

Campbell, Claire E., Devyn L. Cotter, Katherine L. Bottenhorn, Elisabeth Burnor, Hedyeh Ahmadi, W. James Gauderman, Carlos Cardenas-Iniguez, Daniel Hackman, Rob McConnell, Kiros Berhane, Joel Schwartz, Jiu-Chiuan Chen, and Megan M. Herting. 2024. “Air Pollution and Age-Dependent Changes in Emotional Behavior across Early Adolescence in the U.S.” Environmental Research 240:117390. doi:10.1016/j.envres.2023.117390.

Campbell, Claire E., Adam F. Mezher, Sandrah P. Eckel, J. Michael Tyszka, Wolfgang M. Pauli, Bonnie J. Nagel, and Megan M. Herting. 2021. “Restructuring of Amygdala Subregion Apportion across Adolescence.” Developmental Cognitive Neuroscience 48:100883. doi:10.1016/j.dcn.2020.100883.

Campbell, Claire E., Adam F. Mezher, J. Michael Tyszka, Bonnie J. Nagel, Sandrah P. Eckel, and Megan M. Herting. 2022. “Associations between Testosterone, Estradiol, and Androgen Receptor Genotype with Amygdala Subregions in Adolescents.” Psychoneuroendocrinology 137:105604. doi:10.1016/j.psyneuen.2021.105604.

Cao, Lingxiao, Somayeh Maleki Balajoo, Yingxue Gao, Hailong Li, Weijie Bao, Zilin Zhou, Lianqing Zhang, Xinyue Hu, Qiyong Gong, Sarah Genon, and Xiaoqi Huang. 2026. “Common Alterations of Whole and Subregion-Specific Amygdala Intrinsic Functional Connectivity across Psychiatric Disorders: A Meta-Analysis.” Molecular Psychiatry 31(5):2916–26. doi:10.1038/s41380-025-03435-w.

Cardenas-Iniguez, Carlos, Alethea V. de Jesus, Jared Schachner, Shermaine Abad, Joel D. Schwartz, Kirthana Sukumaran, Hedyeh Ahmadi, Daniel A. Hackman, Jiu-Chiuan Chen, and Megan M. Herting. 2025. “Within- and between-Study Site Variations in Ambient Air Pollution Exposure at Ages 9-10 Years in the Adolescent Brain Cognitive Development (ABCD) Study.”

Cardenas-Iniguez, Carlos, Jared N. Schachner, Ka I. Ip, Kathryn E. Schertz, Marybel R. Gonzalez, Shermaine Abad, and Megan M. Herting. 2024. “Building towards an Adolescent Neural Urbanome: Expanding Environmental Measures Using Linked External Data (LED) in the ABCD Study.” Developmental Cognitive Neuroscience 65:101338. doi:10.1016/j.dcn.2023.101338.

Carvalho, Cory, Kalsea Koss, and Niyantri Ravindran. 2026. “The Role of Smartphones in Adolescent-Parent Discrepancy in Reporting Adolescents’ Internalizing Problems.” Development and Psychopathology 38(2):617–29. doi:10.1017/S0954579425100618.

Casey, B. J., Tariq Cannonier, May I. Conley, Alexandra O. Cohen, Deanna M. Barch, Mary M. Heitzeg, Mary E. Soules, Theresa Teslovich, Danielle V. Dellarco, Hugh Garavan, Catherine A. Orr, Tor D. Wager, Marie T. Banich, Nicole K. Speer, Matthew T. Sutherland, Michael C. Riedel, Anthony S. Dick, James M. Bjork, Kathleen M. Thomas, Bader Chaarani, Margie H. Mejia, Donald J. Hagler, M. Daniela Cornejo, Chelsea S. Sicat, Michael P. Harms, Nico U. F. Dosenbach, Monica Rosenberg, Eric Earl, Hauke Bartsch, Richard Watts, Jonathan R. Polimeni, Joshua M. Kuperman, Damien A. Fair, and Anders M. Dale. 2018. “The Adolescent Brain Cognitive Development (ABCD) Study: Imaging Acquisition across 21 Sites.” Developmental Cognitive Neuroscience 32:43–54. doi:10.1016/j.dcn.2018.03.001.

Cattarinussi, Giulia, Yingzhe Zhang, Paola Dazzan, and Divyangana Rakesh. 2026. “The Independent and Joint Effects of Outdoor Air Pollution Exposure and Genetic Risk on Mental Health Trajectories during Adolescence.” medRxiv 2026.07.12.26357864. doi:10.64898/2026.07.12.26357864.

Choi, Jeong-Kyun, Dan Wang, and Aurora P. Jackson. 2019. “Adverse Experiences in Early Childhood and Their Longitudinal Impact on Later Behavioral Problems of Children Living in Poverty.” Child Abuse & Neglect 98:104181. doi:10.1016/j.chiabu.2019.104181.

Clougherty, J. E., J. I. Levy, H. P. Hynes, and J. D. Spengler. 2006. “A Longitudinal Analysis of the Efficacy of Environmental Interventions on Asthma-Related Quality of Life and Symptoms Among Children in Urban Public Housing.” Journal of Asthma 43(5):335–43. doi:10.1080/02770900600701408.

Clougherty, Jane E., and Laura D. Kubzansky. 2009. “A Framework for Examining Social Stress and Susceptibility to Air Pollution in Respiratory Health.” Environmental Health Perspectives 117(9):1351–58. doi:10.1289/ehp.0900612.

deCampo, Danielle M., and Julie L. Fudge. 2012. “Where and What Is the Paralaminar Nucleus? A Review on a Unique and Frequently Overlooked Area of the Primate Amygdala.” Neuroscience & Biobehavioral Reviews 36(1):520–35. doi:10.1016/j.neubiorev.2011.08.007.

Di, Qian, Heresh Amini, Liuhua Shi, Itai Kloog, Rachel Silvern, James Kelly, M. Benjamin Sabath, Christine Choirat, Petros Koutrakis, Alexei Lyapustin, Yujie Wang, Loretta J. Mickley, and Joel Schwartz. 2019. “An Ensemble-Based Model of PM2.5 Concentration across the Contiguous United States with High Spatiotemporal Resolution.” Environment International 130:104909. doi:10.1016/j.envint.2019.104909.

Dolsen, Emily A., Julianna Deardorff, and Allison G. Harvey. 2019. “Salivary Pubertal Hormones, Sleep Disturbance, and an Evening Circadian Preference in Adolescents: Risk Across Health Domains.” Journal of Adolescent Health 64(4):523–29. doi:10.1016/j.jadohealth.2018.10.003.

Dominici, Francesca, Yun Wang, Andrew W. Correia, Majid Ezzati, C. Arden Pope, and Douglas W. Dockery. 2015. “Chemical Composition of Fine Particulate Matter and Life Expectancy.” *Epidemiology (Cambridge*, Mass.) 26(4):556–64. doi:10.1097/EDE.0000000000000297.

Dosanjh, Laura H., Samantha Lauby, Jaime Fuentes, Yessenia Castro, Fiona N. Conway, Frances A. Champagne, Cynthia Franklin, and Bridget Goosby. 2025. “Five Hypothesized Biological Mechanisms Linking Adverse Childhood Experiences with Anxiety, Depression, and PTSD: A Scoping Review.” Neuroscience and Biobehavioral Reviews 171:106062. doi:10.1016/j.neubiorev.2025.106062.

Duffy, Korrina A., Katie A. McLaughlin, and Paige A. Green. 2018. “Early Life Adversity and Health-Risk Behaviors: Proposed Psychological and Neural Mechanisms.” Annals of the New York Academy of Sciences 1428(1):151–69. doi:10.1111/nyas.13928.

Echeverria, Sandra E., Ana V. Diez-Roux, and Bruce G. Link. 2004. “Reliability of Self-Reported Neighborhood Characteristics.” Journal of Urban Health: Bulletin of the New York Academy of Medicine 81(4):682–701. doi:10.1093/jurban/jth151.

Fan, Chun Chieh, Andrew Marshall, Harry Smolker, Marybel R. Gonzalez, Susan F. Tapert, Deanna M. Barch, Elizabeth Sowell, Gayathri J. Dowling, Carlos Cardenas-Iniguez, Jessica Ross, Wesley K. Thompson, and Megan M. Herting. 2021. “Adolescent Brain Cognitive Development (ABCD) Study Linked External Data (LED): Protocol and Practices for Geocoding and Assignment of Environmental Data.” Developmental Cognitive Neuroscience 52:101030. doi:10.1016/j.dcn.2021.101030.

Felitti, Vincent J., MD, FACP, Robert F. Anda MD, MS, Dale Nordenberg MD, David F. Williamson MS, PhD, Alison M. Spitz MS, MPH, Valerie Edwards BA, Mary P. Koss PhD, James S. Marks MD, and MPH. 1998. “Relationship of Childhood Abuse and Household Dysfunction to Many of the Leading Causes of Death in Adults: The Adverse Childhood Experiences (ACE) Study.” American Journal of Preventive Medicine 14(4):245–58. doi:10.1016/S0749-3797(98)00017-8.

Fish, Ari M., Ajay Nadig, Jakob Seidlitz, Paul K. Reardon, Catherine Mankiw, Cassidy L. McDermott, Jonathan D. Blumenthal, Liv S. Clasen, Francois Lalonde, Jason P. Lerch, M. Mallar Chakravarty, Russell T. Shinohara, and Armin Raznahan. 2020. “Sex-Biased Trajectories of Amygdalo-Hippocampal Morphology Change over Human Development.” NeuroImage 204:116122. doi:10.1016/j.neuroimage.2019.116122.

Garavan, H., H. Bartsch, K. Conway, A. Decastro, R. Z. Goldstein, S. Heeringa, T. Jernigan, A. Potter, W. Thompson, and D. Zahs. 2018. “Recruiting the ABCD Sample: Design Considerations and Procedures.” Developmental Cognitive Neuroscience 32:16–22. doi:10.1016/j.dcn.2018.04.004.

Gilpin, Nicholas W., Melissa A. Herman, and Marisa Roberto. 2015. “The Central Amygdala as an Integrative Hub for Anxiety and Alcohol Use Disorders.” Risk Phenotypes for Alcohol and Substance Abuse 77(10):859–69. doi:10.1016/j.biopsych.2014.09.008.

Guxens, Mònica, Małgorzata J. Lubczyńska, Ryan L. Muetzel, Albert Dalmau-Bueno, Vincent W. V. Jaddoe, Gerard Hoek, Aad van der Lugt, Frank C. Verhulst, Tonya White, Bert Brunekreef, Henning Tiemeier, and Hanan El Marroun. 2018. “Air Pollution Exposure During Fetal Life, Brain Morphology, and Cognitive Function in School-Age Children.” Biological Psychiatry 84(4):295–303. doi:10.1016/j.biopsych.2018.01.016.

Hajat, Anjum, Charlene Hsia, and Marie S. O’Neill. 2015. “Socioeconomic Disparities and Air Pollution Exposure: A Global Review.” Current Environmental Health Reports 2(4):440–50. doi:10.1007/s40572-015-0069-5.

Hanson, Jamie L., and Brendon M. Nacewicz. 2021. “Amygdala Allostasis and Early Life Adversity: Considering Excitotoxicity and Inescapability in the Sequelae of Stress.” Frontiers in Human Neuroscience Volume 15-2021. https://www.frontiersin.org/journals/human-neuroscience/articles/10.3389/fnhum.2021.624705.

Herting, Megan M., Cory Johnson, Kathryn L. Mills, Nandita Vijayakumar, Meg Dennison, Chang Liu, Anne-Lise Goddings, Ronald E. Dahl, Elizabeth R. Sowell, Sarah Whittle, Nicholas B. Allen, and Christian K. Tamnes. 2018. “Development of Subcortical Volumes across Adolescence in Males and Females: A Multisample Study of Longitudinal Changes.” NeuroImage 172:194–205. doi:10.1016/j.neuroimage.2018.01.020.

Herzog, Julia I., Janine Thome, Traute Demirakca, Georgia Koppe, Gabriele Ende, Stefanie Lis, Sophie Rausch, Kathlen Priebe, Meike Müller-Engelmann, Regina Steil, Martin Bohus, and Christian Schmahl. 2020. “Influence of Severity of Type and Timing of Retrospectively Reported Childhood Maltreatment on Female Amygdala and Hippocampal Volume.” Scientific Reports 10(1):1903. doi:10.1038/s41598-020-57490-0.

Hobbs, M., E. Moltchanova, L. Marek, K. Yogeeswaran, T. L. Milfont, B. Deng, and C. G. Sibley. 2025. “Environmental Influences on Mental Health: Eight-Year Longitudinal Data Show a Bi-Directional Association between Residential Mobility and Mental Health Outcomes.” Health & Place 94:103487. doi:10.1016/j.healthplace.2025.103487.

Hunsberger, Monica, Jean O’Malley, Torin Block, and Jean C. Norris. 2015. “Relative Validation of Block Kids Food Screener for Dietary Assessment in Children and Adolescents.” Maternal & Child Nutrition 11(2):260–70. doi:10.1111/j.1740-8709.2012.00446.x.

de Jesus, Alethea V., Hedyeh Ahmadi, Daniel A. Hackman, Carlos Cardenas-Iniguez, Jared Schachner, Joel Schwartz, W. James Gauderman, Jiu-Chiuan Chen, and Megan M. Herting. 2025. “Fine Particulate Matter Air Pollution and Longitudinal Gray Matter Development Changes during Early Adolescence: Variation by Neighborhood Disadvantage Level.” Environment International 201:109561. doi:10.1016/j.envint.2025.109561.

Jin, Tingfan, Heresh Amini, Anna Kosheleva, Mahdieh Danesh Yazdi, Yaguang Wei, Edgar Castro, Qian Di, Liuhua Shi, and Joel Schwartz. 2022. “Associations between Long-Term Exposures to Airborne PM2.5 Components and Mortality in Massachusetts: Mixture Analysis Exploration.” Environmental Health 21(1):96. doi:10.1186/s12940-022-00907-2.

Joëls, Marian, and Tallie Z. Baram. 2009. “The Neuro-Symphony of Stress.” Nature Reviews Neuroscience 10(6):459–66. doi:10.1038/nrn2632.

Karcher, Nicole R., and Deanna M. Barch. 2021. “The ABCD Study: Understanding the Development of Risk for Mental and Physical Health Outcomes.” Neuropsychopharmacology 46(1):131–42. doi:10.1038/s41386-020-0736-6.

Keding, Taylor J., and Ryan J. Herringa. 2016. “Paradoxical Prefrontal–Amygdala Recruitment to Angry and Happy Expressions in Pediatric Posttraumatic Stress Disorder.” Neuropsychopharmacology 41(12):2903–12. doi:10.1038/npp.2016.104.

Kessler, Ronald C., Katie A. McLaughlin, Jennifer Greif Green, Michael J. Gruber, Nancy A. Sampson, Alan M. Zaslavsky, Sergio Aguilar-Gaxiola, Ali Obaid Alhamzawi, Jordi Alonso, Matthias Angermeyer, Corina Benjet, Evelyn Bromet, Somnath Chatterji, Giovanni de Girolamo, Koen Demyttenaere, John Fayyad, Silvia Florescu, Gilad Gal, Oye Gureje, Josep Maria Haro, Chi-yi Hu, Elie G. Karam, Norito Kawakami, Sing Lee, Jean-Pierre Lépine, Johan Ormel, José Posada-Villa, Rajesh Sagar, Adley Tsang, T. Bedirhan Üstün, Svetlozar Vassilev, Maria Carmen Viana, and David R. Williams. 2010. “Childhood Adversities and Adult Psychopathology in the WHO World Mental Health Surveys.” British Journal of Psychiatry 197(5):378–85. doi:10.1192/bjp.bp.110.080499.

Kim, Jungmeen, and Dante Cicchetti. 2010. “Longitudinal Pathways Linking Child Maltreatment, Emotion Regulation, Peer Relations, and Psychopathology.” Journal of Child Psychology and Psychiatry 51(6):706–16. doi:10.1111/j.1469-7610.2009.02202.x.

Kim, Mimi S., Shan Luo, Anisa Azad, Claire E. Campbell, Kimberly Felix, Ryan P. Cabeen, Britni R. Belcher, Robert Kim, Monica Serrano-Gonzalez, and Megan M. Herting. 2020. “Prefrontal Cortex and Amygdala Subregion Morphology Are Associated With Obesity and Dietary Self-Control in Children and Adolescents.” Frontiers in Human Neuroscience 14. https://www.frontiersin.org/articles/10.3389/fnhum.2020.563415.

Lahey, Benjamin B., Tyler M. Moore, Antonia N. Kaczkurkin, and David H. Zald. 2021. “Hierarchical Models of Psychopathology: Empirical Support, Implications, and Remaining Issues.” World Psychiatry 20(1):57–63. doi:10.1002/wps.20824.

Lubczyńska, Małgorzata J., Ryan L. Muetzel, Hanan El Marroun, Gerard Hoek, Ingeborg M. Kooter, Errol M. Thomson, Manon Hillegers, Meike W. Vernooij, Tonya White, Henning Tiemeier, and Mònica Guxens. 2021. “Air Pollution Exposure during Pregnancy and Childhood and Brain Morphology in Preadolescents.” Environmental Research 198:110446. doi:10.1016/j.envres.2020.110446.

Margolis, Amy E., Jacob W. Cohen, Bruce Ramphal, Lauren Thomas, Virginia Rauh, Julie Herbstman, and David Pagliaccio. 2022. “Prenatal Exposure to Air Pollution and Early-Life Stress Effects on Hippocampal Subregional Volumes and Associations With Visuospatial Reasoning.” Exposome: Understanding Environmental Impacts on Brain Development and Risk for Psychopathology 2(3):292–300. doi:10.1016/j.bpsgos.2022.05.003.

Mazahir, Fatima A., Ankita Shukla, and Najwa A. Albastaki. 2025. “The Association of Particulate Matter PM2.5 and Nitrogen Oxides from Ambient Air Pollution and Mental Health of Children and Young Adults- a Systematic Review.” Reviews on Environmental Health 40(3):495–536. doi:10.1515/reveh-2024-0120.

McGrath, John J., Ali Al-Hamzawi, Jordi Alonso, Yasmin Altwaijri, Laura H. Andrade, Evelyn J. Bromet, Ronny Bruffaerts, José Miguel Caldas de Almeida, Stephanie Chardoul, Wai Tat Chiu, Louisa Degenhardt, Olga V. Demler, Finola Ferry, Oye Gureje, Josep Maria Haro, Elie G. Karam, Georges Karam, Salma M. Khaled, Viviane Kovess-Masfety, Marta Magno, Maria Elena Medina-Mora, Jacek Moskalewicz, Fernando Navarro-Mateu, Daisuke Nishi, Oleguer Plana-Ripoll, José Posada-Villa, Charlene Rapsey, Nancy A. Sampson, Juan Carlos Stagnaro, Dan J. Stein, Margreet Ten Have, Yolanda Torres, Cristian Vladescu, Peter W. Woodruff, Zahari Zarkov, and Ronald C. Kessler. 2023. “Age of Onset and Cumulative Risk of Mental Disorders: A Cross-National Analysis of Population Surveys from 29 Countries.” The Lancet. Psychiatry 10(9):668–81. doi:10.1016/S2215-0366(23)00193-1.

McLaughlin, Katie A., Jennifer Greif Green, Michael J. Gruber, Nancy A. Sampson, Alan M. Zaslavsky, and Ronald C. Kessler. 2012. “Childhood Adversities and First Onset of Psychiatric Disorders in a National Sample of US Adolescents.” Archives of General Psychiatry 69(11):1151–60. doi:10.1001/archgenpsychiatry.2011.2277.

McLaughlin, Katie A., David Weissman, and Debbie Bitrán. 2019. “Childhood Adversity and Neural Development: A Systematic Review.” Annual Review of Developmental Psychology 1:277–312. doi:10.1146/annurev-devpsych-121318-084950.

Merrick, Melissa T., Derek C. Ford, Katie A. Ports, and Angie S. Guinn. 2018. “Prevalence of Adverse Childhood Experiences From the 2011-2014 Behavioral Risk Factor Surveillance System in 23 States.” JAMA Pediatrics 172(11):1038–44. doi:10.1001/jamapediatrics.2018.2537.

Merz, Emily C., Nim Tottenham, and Kimberly G. Noble. 2018. “Socioeconomic Status, Amygdala Volume, and Internalizing Symptoms in Children and Adolescents.” Journal of Clinical Child & Adolescent Psychology 47(2):312–23. doi:10.1080/15374416.2017.1326122.

Miller, Jonas G., Emily L. Dennis, Sam Heft-Neal, Booil Jo, and Ian H. Gotlib. 2022. “Fine Particulate Air Pollution, Early Life Stress, and Their Interactive Effects on Adolescent Structural Brain Development: A Longitudinal Tensor-Based Morphometry Study.” Cerebral Cortex 32(10):2156–69. doi:10.1093/cercor/bhab346.

Moore, Tyler M., Antonia N. Kaczkurkin, E. Leighton Durham, Hee Jung Jeong, Malerie G. McDowell, Randolph M. Dupont, Brooks Applegate, Jennifer L. Tackett, Carlos Cardenas-Iniguez, Omid Kardan, Gaby N. Akcelik, Andrew J. Stier, Monica D. Rosenberg, Donald Hedeker, Marc G. Berman, and Benjamin B. Lahey. 2020. “Criterion Validity and Relationships between Alternative Hierarchical Dimensional Models of General and Specific Psychopathology.” Journal of Abnormal Psychology 129(7):677–88. doi:10.1037/abn0000601.

Moore, Tyler M., and Benjamin B. Lahey. 2022. “Issues in Estimating Interpretable Lower Order Factors in Second-Order Hierarchical Models: Commentary on Clark et al. (2021).” Clinical Psychological Science 10(3):593–98. doi:10.1177/21677026211035114.

Moore, Tyler M., Elina Visoki, Stirling T. Argabright, Grace E. Didomenico, Ingrid Sotelo, Jeremy D. Wortzel, Areebah Naeem, Ruben C. Gur, Raquel E. Gur, Varun Warrier, Sinan Guloksuz, and Ran Barzilay. 2022. “Modeling Environment through a General Exposome Factor in Two Independent Adolescent Cohorts.” Exposome 2(1):osac010. doi:10.1093/exposome/osac010.

Morrel, Jessica, Michelle Dong, Michael A. Rosario, Devyn L. Cotter, Katherine L. Bottenhorn, and Megan M. Herting. 2025. “A Systematic Review of Air Pollution Exposure and Brain Structure and Function during Development.” Environmental Research 121368. doi:10.1016/j.envres.2025.121368.

Morrel, Jessica, L. Nate Overholtzer, Kirthana Sukumaran, Devyn L. Cotter, Carlos Cardenas-Iniguez, J. Michael Tyszka, Joel Schwartz, Daniel A. Hackman, Jiu-Chiuan Chen, and Megan M. Herting. 2025. “Outdoor Air Pollution Is Related to Amygdala Subregion Volume and Apportionment in Early Adolescence.” Biological Psychiatry Global Open Science 5(5):100544. doi:10.1016/j.bpsgos.2025.100544.

Mujahid, Mahasin S., Ana V. Diez Roux, Jeffrey D. Morenoff, and Trivellore Raghunathan. 2007. “Assessing the Measurement Properties of Neighborhood Scales: From Psychometrics to Ecometrics.” American Journal of Epidemiology 165(8):858–67. doi:10.1093/aje/kwm040.

Nelson, Charles A., Eileen F. Sullivan, and Viviane Valdes. 2025. “Early Adversity Alters Brain Architecture and Increases Susceptibility to Mental Health Disorders.” Nature Reviews Neuroscience 26(10):642–56. doi:10.1038/s41583-025-00948-9.

Nunez, Yanelli, Jaime Benavides, Jenni A. Shearston, Elena M. Krieger, Misbath Daouda, Lucas R. F. Henneman, Erin E. McDuffie, Jeff Goldsmith, Joan A. Casey, and Marianthi-Anna Kioumourtzoglou. 2024. “An Environmental Justice Analysis of Air Pollution Emissions in the United States from 1970 to 2010.” Nature Communications 15(1):268. doi:10.1038/s41467-023-43492-9.

Ojha, Amar, Will Foran, Finnegan J. Calabro, Valerie J. Sydnor, Daniel J. Petrie, Ashley C. Parr, Alyssa Famalette, Natalie Phang, Arshia Sista, Shawn F. Sorrells, and Beatriz Luna. 2025. “Human Amygdala Nuclei Show Distinct Developmental Trajectories from Adolescence to Adulthood in Functional Integration with Prefrontal Circuitry.” Cell Reports 44(9). doi:10.1016/j.celrep.2025.116265.

Orendain, Natalia, Ariana Anderson, Adriana Galván, Susan Bookheimer, and Paul J. Chung. 2023. “A Data-Driven Approach to Categorizing Early Life Adversity Exposure in the ABCD Study.” BMC Medical Research Methodology 23(1):164. doi:10.1186/s12874-023-01983-9.

Oshri, Assaf, Joshua C. Gray, Max M. Owens, Sihong Liu, Erinn Bernstein Duprey, Lawrence H. Sweet, and James MacKillop. 2019. “Adverse Childhood Experiences and Amygdalar Reduction: High-Resolution Segmentation Reveals Associations With Subnuclei and Psychiatric Outcomes.” Child Maltreatment 24(4):400–410. doi:10.1177/1077559519839491.

Oshri, Assaf, Cullin J. Howard, Linhao Zhang, Ava Reck, Zehua Cui, Sihong Liu, Erinn Duprey, Avary I. Evans, Rabeeh Azarmehr, and Charles F. Geier. 2024. “Strengthening through Adversity: The Hormesis Model in Developmental Psychopathology.” Development and Psychopathology 36(5):2390–2406. doi:10.1017/S0954579424000427.

Ousdal, Olga Therese, Anne Marita Milde, Gertrud Sofie Hafstad, Erlend Hodneland, Grete Dyb, Alexander R. Craven, Annika Melinder, Tor Endestad, and Kenneth Hugdahl. 2020. “The Association of PTSD Symptom Severity with Amygdala Nuclei Volumes in Traumatized Youths.” Translational Psychiatry 10(1):288. doi:10.1038/s41398-020-00974-4.

Overholtzer, L. Nate, Carinna Torgerson, Jessica Morrel, Hedyeh Ahmadi, J. Michael Tyszka, and Megan M. Herting. 2025. “Amygdala Subregion Volumes and Apportionment in Preadolescents — Associations with Age, Sex, and Body Mass Index.” Developmental Cognitive Neuroscience 73:101554. doi:10.1016/j.dcn.2025.101554.

Page, Chloe E., Sean W. Biagiotti, Pia J. Alderman, and Shawn F. Sorrells. 2022. “Immature Excitatory Neurons in the Amygdala Come of Age during Puberty.” Developmental Cognitive Neuroscience 56:101133. doi:10.1016/j.dcn.2022.101133.

Pagliaccio, David, Julie B. Herbstman, Frederica Perera, Deliang Tang, Jeff Goldsmith, Bradley S. Peterson, Virginia Rauh, and Amy E. Margolis. 2020. “Prenatal Exposure to Polycyclic Aromatic Hydrocarbons Modifies the Effects of Early Life Stress on Attention and Thought Problems in Late Childhood.” Journal of Child Psychology and Psychiatry 61(11):1253–65. doi:10.1111/jcpp.13189.

Pagliaccio, David, Kate T. Tran, Elina Visoki, Grace E. DiDomenico, Randy P. Auerbach, and Ran Barzilay. 2024. “Probing the Digital Exposome: Associations of Social Media Use Patterns with Youth Mental Health.” NPP—Digital Psychiatry and Neuroscience 2(1):5. doi:10.1038/s44277-024-00006-9.

Palmer, Clare E., Diliana Pecheva, John R. Iversen, Donald J. Hagler, Leo Sugrue, Pierre Nedelec, Chun Chieh Fan, Wesley K. Thompson, Terry L. Jernigan, and Anders M. Dale. 2022. “Microstructural Development from 9 to 14 Years: Evidence from the ABCD Study.” Developmental Cognitive Neuroscience 53:101044. doi:10.1016/j.dcn.2021.101044.

Pauli, Wolfgang M., Amanda N. Nili, and J. Michael Tyszka. 2018. “A High-Resolution Probabilistic in Vivo Atlas of Human Subcortical Brain Nuclei.” Scientific Data 5(1):180063. doi:10.1038/sdata.2018.63.

Paus, Tomáš, Matcheri Keshavan, and Jay N. Giedd. 2008. “Why Do Many Psychiatric Disorders Emerge during Adolescence?” Nature Reviews Neuroscience 9(12):947–57. doi:10.1038/nrn2513.

Peverill, Matthew, Maya L. Rosen, Lucy A. Lurie, Kelly A. Sambrook, Margaret A. Sheridan, and Katie A. McLaughlin. 2023. “Childhood Trauma and Brain Structure in Children and Adolescents.” Developmental Cognitive Neuroscience 59:101180. doi:10.1016/j.dcn.2022.101180.

Pfeifer, Jennifer H., and Nicholas B. Allen. 2021. “Puberty Initiates Cascading Relationships Between Neurodevelopmental, Social, and Internalizing Processes Across Adolescence.” Adolescent Brain Development and Psychopathology 89(2):99–108. doi:10.1016/j.biopsych.2020.09.002.

Phelps, Elizabeth A., and Joseph E. LeDoux. 2005. “Contributions of the Amygdala to Emotion Processing: From Animal Models to Human Behavior.” Neuron 48(2):175–87. doi:10.1016/j.neuron.2005.09.025.

Prager, Eric M., Hadley C. Bergstrom, Gary H. Wynn, and Maria F. M. Braga. 2016. “The Basolateral Amygdala γ-Aminobutyric Acidergic System in Health and Disease.” Journal of Neuroscience Research 94(6):548–67. doi:10.1002/jnr.23690.

R Core Team. 2025. “R: A Language and Environment for Statistical Computing (Version 3.2. 3)[Computer Software]. R Foundation for Statistical Computing, Vienna, Austria.”

Rakesh, Divyangana, Andrew Zalesky, and Sarah Whittle. 2022. “Assessment of Parent Income and Education, Neighborhood Disadvantage, and Child Brain Structure.” JAMA Network Open 5(8):e2226208. doi:10.1001/jamanetworkopen.2022.26208.

Renzetti, Stefano, Paul Curtin, Allan C. Just, Ghalib Bello, and Chris Gennings. 2021. “gWQS: Generalized Weighted Quantile Sum Regression.”

Requia, Weeberb J., Qian Di, Rachel Silvern, James T. Kelly, Petros Koutrakis, Loretta J. Mickley, Melissa P. Sulprizio, Heresh Amini, Liuhua Shi, and Joel Schwartz. 2020. “An Ensemble Learning Approach for Estimating High Spatiotemporal Resolution of Ground-Level Ozone in the Contiguous United States.” Environmental Science & Technology 54(18):11037–47. doi:10.1021/acs.est.0c01791.

Reuben, Aaron, Louise Arseneault, Andrew Beddows, Sean D. Beevers, Terrie E. Moffitt, Antony Ambler, Rachel M. Latham, Joanne B. Newbury, Candice L. Odgers, Jonathan D. Schaefer, and Helen L. Fisher. 2021. “Association of Air Pollution Exposure in Childhood and Adolescence With Psychopathology at the Transition to Adulthood.” JAMA Network Open 4(4):e217508. doi:10.1001/jamanetworkopen.2021.7508.

Roozendaal, Benno, Bruce S. McEwen, and Sumantra Chattarji. 2009. “Stress, Memory and the Amygdala.” Nature Reviews Neuroscience 10(6):423–33. doi:10.1038/nrn2651.

de la Rosa, Rosemarie, Austin Le, Stephanie Holm, Morgan Ye, Nicole R. Bush, Danielle Hessler, Kadiatou Koita, Monica Bucci, Dayna Long, and Neeta Thakur. 2024. “Associations Between Early-Life Adversity, Ambient Air Pollution, and Telomere Length in Children.” Psychosomatic Medicine 86(5):422–30. doi:10.1097/PSY.0000000000001276.

Rosario, Michael A., Kirthana Sukumaran, Katherine L. Bottenhorn, Alethea de Jesus, Carlos Cadenas-Iniguez, Hedyeh Ahmadi, Rima Habre, Shermaine Abad, Jacob G. Pine, Deanna M. Barch, Joel Schwartz, Daniel A. Hackman, Jiu-Chiuan Chen, and Megan M. Herting. 2026. “Ambient Pollution Components and Sources Are Associated with Hippocampal Architecture and Memory in Pre-Adolescents.” BMC Medicine. doi:10.1186/s12916-026-04950-5.

Russell, Justin D., Monica A. Marsee, and Carl F. Weems. 2021. “Developmental Variation in Amygdala Volumes: Modeling Differences Across Time, Age, and Puberty.” Imaging Biomarkers and Outcome Prediction 6(1):117–25. doi:10.1016/j.bpsc.2020.08.006.

Sambuco, Nicola, Margaret M. Bradley, and Peter J. Lang. 2023. “Hippocampal and Amygdala Volumes Vary with Transdiagnostic Psychopathological Dimensions of Distress, Anxious Arousal, and Trauma.” Biological Psychology 177:108501. doi:10.1016/j.biopsycho.2023.108501.

Sangkham, Sarawut. 2026. “PM2.5 Is a Toxic Mixture: Not Just a Matter of Concentration.” Chemical Research in Toxicology 39(3):249–52. doi:10.1021/acs.chemrestox.5c00455.

Sepahvand, Tayebeh, Kyron D. Power, Tian Qin, and Qi Yuan. 2023. “The Basolateral Amygdala: The Core of a Network for Threat Conditioning, Extinction, and Second-Order Threat Conditioning.” Biology 12(10):1274. doi:10.3390/biology12101274.

Sharp, B. M. 2017. “Basolateral Amygdala and Stress-Induced Hyperexcitability Affect Motivated Behaviors and Addiction.” Translational Psychiatry 7(8):e1194–e1194. doi:10.1038/tp.2017.161.

Shin, Lisa M., Scott L. Rauch, and Roger K. Pitman. 2006. “Amygdala, Medial Prefrontal Cortex, and Hippocampal Function in PTSD.” Annals of the New York Academy of Sciences 1071(1):67–79. doi:10.1196/annals.1364.007.

Smolker, Harry R., Colleen E. Reid, Naomi P. Friedman, and Marie T. Banich. 2024. “The Association between Exposure to Fine Particulate Air Pollution and the Trajectory of Internalizing and Externalizing Behaviors during Late Childhood and Early Adolescence: Evidence from the Adolescent Brain Cognitive Development (ABCD) Study.” Environmental Health Perspectives 132(8):087001. doi:10.1289/EHP13427.

Sorrells, Shawn F., Mercedes F. Paredes, Arantxa Cebrian-Silla, Kadellyn Sandoval, Dashi Qi, Kevin W. Kelley, David James, Simone Mayer, Julia Chang, Kurtis I. Auguste, Edward F. Chang, Antonio J. Gutierrez, Arnold R. Kriegstein, Gary W. Mathern, Michael C. Oldham, Eric J. Huang, Jose Manuel Garcia-Verdugo, Zhengang Yang, and Arturo Alvarez-Buylla. 2018. “Human Hippocampal Neurogenesis Drops Sharply in Children to Undetectable Levels in Adults.” Nature 555(7696):377–81. doi:10.1038/nature25975.

Sukumaran, Kirthana, Katherine L. Bottenhorn, Michael A. Rosario, Carlos Cardenas-Iniguez, Rima Habre, Shermaine Abad, Joel Schwartz, Daniel A. Hackman, J. C. Chen, and Megan M. Herting. 2025. “Sources and Components of Fine Air Pollution Exposure and Brain Morphology in Preadolescents.” Science of The Total Environment 979:179448. doi:10.1016/j.scitotenv.2025.179448.

Sukumaran, Kirthana, Katherine L. Botternhorn, Joel Schwartz, Jim Gauderman, Carlos Cardenas-Iniguez, Rob McConnell, Daniel A. Hackman, Kiros Berhane, Hedyeh Ahmadi, Shermaine Abad, Rima Habre, and Megan M. Herting. 2024. “Associations between Fine Particulate Matter Components, Their Sources, and Cognitive Outcomes in Children Ages 9–10 Years Old from the United States.” Environmental Health Perspectives 132(10):107009. doi:10.1289/EHP14418.

Swedo, Elizabeth A., Phyllis Holditch Niolon, Kayla N. Anderson, Jingjing Li, Nancy Brener, Jonetta Mpofu, Maria V. Aslam, and J. Michael Underwood. 2024. “Prevalence of Adverse Childhood Experiences Among Adolescents.” Pediatrics 154(5):e2024066633. doi:10.1542/peds.2024-066633.

Tyszka, J. Michael, and Wolfgang M. Pauli. 2016. “In Vivo Delineation of Subdivisions of the Human Amygdaloid Complex in a High-Resolution Group Template.” Human Brain Mapping 37(11):3979–98. doi:10.1002/hbm.23289.

Vachon, David D., Robert F. Krueger, Fred A. Rogosch, and Dante Cicchetti. 2015. “Assessment of the Harmful Psychiatric and Behavioral Effects of Different Forms of Child Maltreatment.” JAMA Psychiatry 72(11):1135–42. doi:10.1001/jamapsychiatry.2015.1792.

VanTieghem, Michelle, Marta Korom, Jessica Flannery, Tricia Choy, Christina Caldera, Kathryn L. Humphreys, Laurel Gabard-Durnam, Bonnie Goff, Dylan G. Gee, Eva H. Telzer, Mor Shapiro, Jennifer Y. Louie, Dominic S. Fareri, Niall Bolger, and Nim Tottenham. 2021. “Longitudinal Changes in Amygdala, Hippocampus and Cortisol Development Following Early Caregiving Adversity.” Developmental Cognitive Neuroscience 48:100916. doi:10.1016/j.dcn.2021.100916.

Vargas, Teresa G., Katie A. McLaughlin, and Divyangana Rakesh. 2025. “Testing Moderators for Associations of Neighborhood Adversity With Psychopathology and Cognitive Outcomes.” Developmental Science 28(5):e70055. doi:10.1111/desc.70055.

Volkow, Nora D., George F. Koob, Robert T. Croyle, Diana W. Bianchi, Joshua A. Gordon, Walter J. Koroshetz, Eliseo J. Pérez-Stable, William T. Riley, Michele H. Bloch, Kevin Conway, Bethany G. Deeds, Gayathri J. Dowling, Steven Grant, Katia D. Howlett, John A. Matochik, Glen D. Morgan, Margaret M. Murray, Antonio Noronha, Catherine Y. Spong, Eric M. Wargo, Kenneth R. Warren, and Susan R. B. Weiss. 2018. “The Conception of the ABCD Study: From Substance Use to a Broad NIH Collaboration.” Developmental Cognitive Neuroscience 32:4–7. doi:10.1016/j.dcn.2017.10.002.

Vyas, Chirag M., Macarius Donneyong, David Mischoulon, Grace Chang, Heike Gibson, Nancy R. Cook, JoAnn E. Manson, Charles F. Reynolds, and Olivia I. Okereke. 2020. “Associations between Race and Ethnicity and Late-Life Depression Severity, Symptom Burden and Care.” JAMA Network Open 3(3):e201606. doi:10.1001/jamanetworkopen.2020.1606.

Wade, Mark, Liam Wright, and Katherine E. Finegold. 2022. “The Effects of Early Life Adversity on Children’s Mental Health and Cognitive Functioning.” Translational Psychiatry 12(1):244. doi:10.1038/s41398-022-02001-0.

Waters, Renée C., and Elizabeth Gould. 2022. “Early Life Adversity and Neuropsychiatric Disease: Differential Outcomes and Translational Relevance of Rodent Models.” Frontiers in Systems Neuroscience 16. doi:10.3389/fnsys.2022.860847.

Wierenga, Lara, Marieke Langen, Sara Ambrosino, Sarai van Dijk, Bob Oranje, and Sarah Durston. 2014. “Typical Development of Basal Ganglia, Hippocampus, Amygdala and Cerebellum from Age 7 to 24.” NeuroImage 96:67–72. doi:10.1016/j.neuroimage.2014.03.072.

Wodtke, Geoffrey T., Kerry Ard, Clair Bullock, Kailey White, and Betsy Priem. 2022. “Concentrated Poverty, Ambient Air Pollution, and Child Cognitive Development.” Science Advances 8(48):eadd0285. doi:10.1126/sciadv.add0285.

Wu, Pengpeng, Qian Guo, Yuchen Zhao, Mengyao Bian, Suzhen Cao, Junfeng (Jim) Zhang, and Xiaoli Duan. 2024. “Emerging Concern on Air Pollution and Health: Trade-off between Air Pollution Exposure and Physical Activity.” Eco-Environment & Health 3(2):202–7. doi:10.1016/j.eehl.2024.01.012.

Zundel, Clara G., Patrick Ryan, Cole Brokamp, Autumm Heeter, Yaoxian Huang, Jeffrey R. Strawn, and Hilary A. Marusak. 2022. “Air Pollution, Depressive and Anxiety Disorders, and Brain Effects: A Systematic Review.” Neurotoxicology 93:272–300. doi:10.1016/j.neuro.2022.10.011.

